# Tau, XMAP215/Msps and Eb1 co-operate interdependently to regulate microtubule polymerisation and bundle formation in axons

**DOI:** 10.1101/2020.08.19.257808

**Authors:** Ines Hahn, Andre Voelzmann, Jill Parkin, Judith Fuelle, Paula G Slater, Laura A Lowery, Natalia Sanchez-Soriano, Andreas Prokop

## Abstract

The formation and maintenance of microtubules requires their polymerisation, but little is known about how this polymerisation is regulated in cells. Focussing on the essential microtubule bundles in axons of *Drosophila* and *Xenopus* neurons, we show that the plus-end scaffold Eb1, the polymerase XMAP215/Msps and the lattice-binder Tau co-operate interdependently to promote microtubule polymerisation and bundle organisation during axon development and maintenance. Eb1 and XMAP215/Msps promote each other’s localisation at polymerising microtubule plus-ends. Tau outcompetes Eb1-binding along microtubule lattices, thus preventing depletion of Eb1 tip pools. The three factors genetically interact and show shared mutant phenotypes: reductions in axon growth, comet size, comet number and comet velocity, as well as prominent deterioration of parallel microtubule bundles into disorganised curled conformations. This microtubule curling is caused by Eb1 plus-end depletion which impairs spectraplakin-mediated guidance of extending microtubules into parallel bundles. Our demonstration that Eb1, XMAP215/Msps and Tau co-operate during the regulation of microtubule polymerisation and bundle organisation, offers new conceptual explanations for developmental and degenerative axon pathologies and how to treat them.

**Summary statement:** Eb1, XMAP215 and tau co-operate interdependently in axons to promote the polymerisation of microtubules and their organisation into the parallel bundles required for axonal transport.

## Introduction

Axons are the enormously long cable-like cellular processes of neurons that wire nervous systems. In humans, axons of ≤15µm diameter can be up to two meters long (Prokop, 2020; Prokop, 2021). They are constantly exposed to mechanical challenges, yet have to survive for up to a century; we lose ∼40% of axons towards high age and far more in neurodegenerative diseases (Adalbert & Coleman, 2012; Calkins, 2013; Marner *et al*, 2003).

Their growth and maintenance absolutely require parallel bundles of microtubules (MTs) that run all along axons, providing the highways for life-sustaining transport and driving morphogenetic events. Consequently, bundle decay through MT loss or curling is a common feature in axon pathologies (summarised in (Hahn *et al*, 2019; Prokop, 2021). Key roles must be played by MT polymerisation, which is not only essential for the *de novo* formation of MT bundles occurring during axon growth in development, plasticity or regeneration, but also to repair damaged and replace senescent MTs during long-term maintenance (Gasic & Mitchison, 2019; Schaedel *et al*, 2015; Voelzmann *et al*, 2016). However, the molecular mechanisms regulating MT polymerisation in axons are surprisingly little understood.

MT polymerisation is primarily understood *in vitro*, where MTs can undergo polymerisation in the presence of nucleation seeds and tubulin heterodimers; the addition of catalytic factors such as CLASPs, stathmins, tau, Eb proteins or XMAP215 can enhance and refine the process (Aher *et al*, 2020; Al-Bassam *et al*, 2010; Brouhard *et al*, 2008; Drechsel *et al*, 1992; Li *et al*, 2012; Manna *et al*, 2009; Zanic *et al*, 2013). However, we do not know whether mechanisms observed in reconstitution assays are biologically relevant in the context of axons (Voelzmann *et al*., 2016), especially when considering that none of the above-mentioned factors has genetic links to human neurological disorders on OMIM® (Online Mendelian Inheritance in Man), except Tau/MAPT which executes numerous disease-relevant functions also when detached from MTs (Morris *et al*, 2011).

To identify relevant factors regulating axonal MT polymerisation, we use *Drosophila* primary neurons as one consistent model, which is amenable to combinatorial genetics as a powerful strategy to decipher complex regulatory networks (Prokop *et al*, 2013). Our previous loss-of-function studies of 9 MT plus-end-associating factors in these *Drosophila* neurons (CLASP, CLIP190, dynein heavy chain, APC, p150^Glued^, Eb1, Short stop/Shot, doublecortin, Lis1) have taken axon length as a crude proxy readout for net polymerisation, mostly revealing relatively mild axon length phenotypes, with the exception of Eb1 and Shot which cause severe axon shortening {Beaven, 2015 #6487; Sánchez-Soriano, 2009 #7139; Alves-Silva, 2012 #4930; A.V., unpublished data}.

Here we have incorporated more candidate factors and readouts to take these analyses to the next level. We show that three factors, Eb1, XMAP215/Msps and Tau, share a unique combination of mutant phenotypes in culture and *in vivo*, including reduced axonal MT polymerisation in frog and fly neurons. Our data reveal that the three factors co-operate. Eb1 and XMAP215/Msps act interdependently at MT plus-ends, whereas Tau acts through a novel mechanism: it outcompetes Eb1-binding along MT lattices, thus preventing the depletion of Eb1 pools at polymerising MT plus-ends. By upholding these Eb1 pools, the functional trio also promotes the bundle conformation of axonal MTs through a guidance mechanism mediated by the spectraplakin Shot. Our work uniquely integrates molecular mechanisms into understanding of MT regulation that is biologically relevant for axon growth, maintenance and disease.

## Methods

### Fly stocks

Loss-of-function mutant stocks used in this study were the deficiencies *Df(3R)Antp17* (*tub^def^*; removing both *αtub84B* and *αtub84D*; (Duncan & Kaufman, 1975; Jenkins *et al*, 2017), *Df(2L)Exel6015* (*stai^Df^*; (Duncan *et al*, 2013), *Df(3L)BSC553* (*clasp^Df^*; Bloomington stock #25116; Beaven *et al*, 2015, *Df(3R)tauMR22* (*tau^Df^*; Doerflinger *et al*, 2003) and the loss-of-function mutant alleles *α-tub84B^KO^* (an engineered null-allele; Jenkins *et al*., 2017), *chromosome bows^2^* (*clasp^2^*, an amorph allele; Inoue *et al*, 2000), *Eb1^04524^*, *Eb1^5^* (strong loss-of-function mutant alleles; Elliott *et al*, 2005), *futsch^P158^* (*MAP1B^-^*; a deficiency uncovering the *futsch* locus; Hummel *et al*, 2000), *msps^A^* (a small deletion causing a premature stop after 370 amino acids; gift from H. Ohkura, unpublished), *msps^146^* (Brittle & Ohkura, 2005), *sentin^ΔB^* (*short spindle2^ΔB^, ssp2^ΔB^*; Gluszek *et al*, 2015), *tacc^1^* (*dTACC^1^*; Gergely *et al*, 2000), *shot^3^* (the strongest available allele of *short stop*; Kolodziej *et al*, 1995; Sánchez-Soriano *et al*, 2009), *stai^KO^* (Yang *et al*, 2012), *tau^KO^* (a null allele; Burnouf *et al*, 2016). Gal4 driver lines used were *elav-Gal4* (1^st^ and 3^rd^ chromosomal, both expressing pan-neuronally at all stages; Luo *et al*, 1994), *GMR31F10-Gal4* (Bloomington #49685; expressing in T1 medulla neurons; Qu *et al*, 2019). Lines for targeted gene expression were *UAS-Eb1-GFP* and *UAS-shot-Ctail-GFP* (Alves-Silva *et al*, 2012), *UAS-shot^ΔABD^-GFP* (Qu, 2015), *UAS-shot^3MTLS^-GFP* (Alves-Silva *et al*., 2012), *UAS-dtau-GFP* (Doerflinger *et al*., 2003), *UAS-GFP-α-tubulin84B* (Grieder *et al*, 2000) and further lines generated here (see below).

### *Drosophila* primary cell culture

*Drosophila* primary neuron cultures were done as described previously (Prokop *et al*, 2012; Qu *et al*, 2017). Stage 11 embryos were treated for 90 s with bleach to remove the chorion, sterilized for ∼30 s in 70% ethanol, washed in sterile Schneider’s medium containing 20% fetal calf serum (Schneider’s/FCS; Gibco), and eventually homogenized with micro-pestles in 1.5 centrifuge tubes containing 21 embryos per 100 μl dispersion medium (Prokop *et al*., 2012) and left to incubated for 4 min at 37°C. Dispersion was stopped with 200 μl Schneider’s/FCS, cells were spun down for 4 mins at 650 g, supernatant was removed and cells re-suspended in 90 µl of Schneider’s/FCS, and 30 μl drops were placed in culture chambers and covered with cover slips. Cells were allowed to adhere to cover slips for 90-120 min either directly on glass or on cover slips coated with a 5 µg/ml solution of concanavalin A, and then grown as a hanging drop culture at 26°C as indicated.

To eliminate a potential maternal rescue of mutants (i.e. reduction of the mutant phenotype due to normal gene product deposition from the wild-type gene copy of the heterozygous mothers in oocytes (Prokop, 2013), we used a pre-culture strategy (Prokop *et al*., 2012; Sánchez-Soriano *et al*, 2010) where cells were incubated in a tube for 5-7 days before they were plated on coverslips.

For cultures from larval brains, L3 brains (2-3 per cover slip) were dissected in PBS, transferred into Schneider’s/FCS medium, washed three times with medium and then proceed with homogenisation and dispersion as explained above

Transfection of *Drosophila* primary neurons was executed as described previously (Qu *et al*., 2019). In brief, 70-75 embryos per 100 μl dispersion medium were used. After the washing step and centrifugation, cells were re-suspended in 100 μl transfection medium [final media containing 0.1-0.5 μg DNA and 2 μl Lipofecatmine 2000 (L2000, Invitrogen), incubation following manufacturer’s protocols (Thermo Fisher, Invitrogen) and kept for 24 hrs at 26°C. Cells were then treated again with dispersion medium, re-suspended in culture medium and plated out as described above.

### *Xenopus* primary neuron experiments

Xenopus primary neuron cultures were obtained from embryonic neural tube. Eggs collected from female X. laevis frogs were fertilised in vitro, dejellied and cultured following standard methods (Sive *et al*, 2010). Embryos were grown to stage 22–24 (Nieuwkoop & J., 1994), and neural tubes were dissected as described (Lowery *et al*, 2012). Three neural tubes were transferred to an Eppendorf tube containing 150 uL CMF-MMR (0.1 M NaCl, 2.0 mM KCl, 1.0 mM EDTA, 5.0 mM HEPES, pH 7.4), 10 min later centrifuged at 1000 g for 5 min, and 150 uL of Steinberg’s solution (58 mM NaCl, 0.67 mM KCl, 0.44 mM Ca(NO_3_)_2_, 1.3 mM MgSO_4_, 4.6 mM Tris, pH 7.8) was added to the supernatant to follow with the tissue dissociation using a fire polished glass Pasteur pipet. Cells were seeded in 100 ug/mL Poly-L-lysine and 10 μg/ml laminin pre-treated 60mm plates, and after 2 hr the media was replaced by plating culture media (50% Ringer’s, 49% L-15 media, 1% Fetal Bovine Serum, 25 ng/μl NT3 and BDNF, plus 50 μg/ml penicillin/streptomycin and gentamycin, pH 7.4 and filter sterilized) and kept for 24 hr before imaging.

All experiments were approved by the Boston College Institutional Animal Care and Use Committee and performed according to national regulatory standards.

The embryos were injected four times in dorsal blastomeres at two-to-four cell stage with 6 ng of the validated XMAP215 morpholino (MO; Lowery *et al*, 2013), 10 ng of the validated Tau MO (Liu *et al*, 2015), and/or 5 ng of a newly designed splice site MO for EB3 (3’CTCCCAATTGTCACCTACTTTGTCG5’; for verification see Fig3-S1), in order to obtain a 50% knockdown of each.

To assess EB1 comet dynamics and comet amounts, 300 pg of MACF43-Ctail::GFP, an Eb protein-binding 43-residue fragment derived from the C-terminal regions of hMACF2 (human microtubule actin crosslinking factor 2; Fig.3F-F’’’; Honnappa *et al*, 2009; Slater *et al*, 2019), was co-injected with the MO.

### Immunohistochemistry

Primary fly neurons were fixed in 4% paraformaldehyde (PFA; in 0.05 M phosphate buffer, pH 7–7.2) for 30 min at room temperature (RT). For anti-Eb1 and anti-GTP-tubulin staining, cells were fixed for 10 mins at −20°C in +TIP fix (90% methanol, 3% formaldehyde, 5 mM sodium carbonate, pH 9; stored at −80°C and added to the cells) (Rogers *et al*, 2002), then washed in PBT (PBS with 0.3% TritonX). Antibody staining and washes were performed with PBT. Staining reagents: anti-tubulin (clone DM1A, mouse, 1:1000, Sigma; alternatively, clone YL1/2, rat, 1:500, Millipore Bioscience Research Reagents); anti-DmEb1 (gift from H. Ohkura; rabbit, 1:2000; (Elliott *et al*., 2005); anti-GTP-tubulin (hMB11; human, 1:200; AdipoGen; Dimitrov *et al*, 2008); anti-Shot (1:200, guinea pig; Strumpf & Volk, 1998); anti-Elav (Elav-7E8A10; rat, 1:1000; Developmental Studies Hybridoma Bank, The University of Iowa, IA, USA) (O’Neill *et al*, 1994); anti-GFP (ab290, Abcam, 1:500); Cy3-conjugated anti-HRP (goat, 1:100, Jackson ImmunoResearch); F-actin was stained with phalloidin conjugated with TRITC/Alexa647, FITC or Atto647N (1:100 or 1:500; Invitrogen and Sigma). Specimens were embedded in ProLong Gold Antifade Mountant (ThermoFisher Scientific).

### Western blot analysis of *Xenopus* embryos

For protein extraction, 10 embryos were transferred to a centrifuge tube with 800 µl lysis buffer (50 mM Tris pH 7.5, 5% glycerol, 0.2% IGEPAL/NP-40, 1 mM EDTA, 1.5 mM MgCl_2_, 125 mM NaCl, 25 mM NaF, 1 mM Na_3_VO_4_), homogenised with a sterile pestle and, after 10 min, centrifuged at 13000 rpm for 15-20 min. The supernant was collected and the protein concentration determined with the Micro BCA^TM^ Protein Assay Kit (Thermo Fisher Scientific). 80 µg protein were loaded into a 10 % SDS gel and stained with anti-Tau (clone Tau46, T9450, mouse, 1:1000, Sigma-Aldrich).

### Dissection of adult heads

For *in vivo* studies, brain dissections were performed in Dulbecco’s PBS (Sigma, RNBF2227) after briefly sedating them on ice. Dissected brains with their laminas and eyes attached were placed into a drop of Dulbecco’s PBS on MatTek glass bottom dishes (P35G1.5-14C), covered by coverslips and immediately imaged with a 3i Marianas Spinning Disk Confocal Microscope.

### Microscopy and data analysis

Standard imaging was performed with AxioCam 506 monochrome (Carl Zeiss Ltd.) or MatrixVision mvBlueFox3-M2 2124G digital cameras mounted on BX50WI or BX51 Olympus compound fluorescent microscopes. For the analysis of *Drosophila* and *Xenopus* primary neurons, we used the following parameters:

Axon length was measured from cell body to growth cone tip using the segmented line tool of ImageJ (Alves-Silva *et al*., 2012; Sánchez-Soriano *et al*., 2010).

Degree of disorganised MT curling in axons was measured as “MT disorganisation index” (MDI) described previously (Qu *et al*., 2019; Qu *et al*., 2017); in short: the area of disorganised curling was measured with the freehand selection in ImageJ; this value was then divided by axon length (see above) multiplied by 0.5 μm (typical axon diameter, thus approximating the expected area of the axon if it were properly bundled).

Eb1 comet amounts were approximated by using product of comet mean intensity and length. For this, a line was drawn through each comet (using the segmented line tool in FIJI) and length as well as mean staining intensity (of Eb1 or GTP-tub in fixed *Drosophila* and MACF43::GFP in a movie still in *Xenopus* neurons) was determined.

To measure MT curling in the optic lobe of adult flies, *GMR31F10-Gal4* (Bloomington #49685) was used to express *UAS-α-tubulin84B-GFP* (Grieder *et al*., 2000) in a subset of lamina axons which projects within well-ordered medulla columns (Prokop & Meinertzhagen, 2006). Flies were left to age for 26-27 days (about half their life expectancy) and then their brains were dissected out, mounted in Mattek dishes and imaged using a 3i spinning disk confocal system at the ITM Biomedecial imaging facility at the University of Liverpool. A section of the medulla columns comprising the 4 most proximal axonal terminals was used to quantify the number of swellings and regions with disorganised curled MTs.

To measure MT polymerisation dynamics, movies were collected on an Andor Dragonfly200 spinning disk upright confocal microscope (with a Leica DM6 FS microscope frame) and using a *100x/1.40 UPlan SAPO (Oil)* objective. Samples where excited using 488nm (100%) and 561nm (100%) diode lasers via Leica GFP and RFP filters respectively. Images where collected using a Zyla 4.2 Plus sCMOS camera with a camera gain of 1x. The incubation temperature was set to 26°C. Time lapse movies were constructed from images taken every 1 s for 1 mins. To measure comet velocity and lifetime, a line was drawn that followed the axon using the segmented line tool in ImageJ. A kymograph was then constructed from average intensity in FIJI using the KymoResliceWide macro (Cell Biology group, Utrecht University) and events scored via the Velocity Measurement Tool Macro (Volker Baecker, INSERM, Montpellier, RIO Imaging; J. Rietdorf, FMI Basel; A. Seitz, EMBL Heidelberg). For each condition at least 15 cells were analysed in ≥2 independent repeats.

Time lapse imaging for *Xenopus* primary cultures was performed with a CSU-X1M 5000 spinning-disk confocal (Yokogawa, Tokyo, Japan) on a Zeiss Axio Observer inverted motorized microscope with a Zeiss 63× Plan Apo 1.4 numerical aperture lens (Zeiss, Thornwood, NY). Images were acquired with an ORCA R2 charge-coupled device camera (Hamamatsu, Hamamatsu, Japan) controlled with Zen software. Time lapse movies were constructed from images taken every 2 s for 1 min. The MACF43 comets’ velocities and lifetime were analysed with plusTipTracker software. The same parameters were used for all movies: maximum gap length, eight frames; minimum track length, three frames; search radius range, 5–12 pixels; maximum forward angle, 50°; maximum backward angle, 10°; maximum shrinkage factor, 0.8; fluctuation radius, 2.5 pixels; and time interval 2 s.

Images were derived from at least 3 independent experimental repeats performed on different days, for each of which at least 2 independent culture wells were analysed by taking a minimum of 20 images per slide. For statistical analyses, Kruskal–Wallis one-way ANOVA with *post hoc* Dunn’s test or Mann–Whitney Rank Sum Tests (indicated as P_MW_) were used to compare groups, r and p-value for correlation were determined via non-parametric Spearman correlation analysis (tests showed that data are not distributed normally). All raw data of our analyses are provided as supplementary Excel files T1-6.

### Molecular biology

To generate the *UAS-msps^FL^-GFP* (aa1-2050) and *UAS-msps^ΔCterm^* (aa1-1322) constructs, eGFP was PCR-amplified from pcDNA3-EGFP and *msps* sequences from cDNA clone *LP04448* (DGRC; FBcl0189229) using the following primers:

*msps^FL^* and *msps^ΔCterm^* fw: GAATAGGGAATTGGGAATTCGTTAGGCGCGCCAACATGGCCGAGGACACAGAGTAC

*msps^FL^* rev: *CAAGAAAGAGAATCATGCCCAAGGGCCCGGTAGCGGCAGCGGTAGCGTGAGCAAGGGC GAG*

*msps^ΔCterm^* rev: *GATGGAGGGTCTAAAATCGCATATGGGTAGCGGCAGCGGTAGCGTGAGCAAGGGCGAG GAG*

*eGFP* fw: *GAGAATCATGCCCAAGGGCCCGGTAGCGGCAGCGGTAGCGTGAGCAAGGGCGAGGAG CTG*

*eGFP* rev: *CTCTCGGCATGGACGAGCTGTACAAGTAGGCGGCCGCCTCGAGGGTACCTCTAGAG*

The *msps* and *eGFP* sequences were introduced into *pUAST-attB* via Gibson assembly (ThermoFisher) using *EcoRI* and *XhoI*. To generate transgenic fly lines, *pUAST-attB* constructs were integrated into *PBac{yellow[+]-attP-9A}VK00024* (Bloomington line #9742) via PhiC31-mediated recombination (outsourced to Bestgene Inc).

## Results

### Eb1, Msps/XMAP215 and Tau share the same combination of loss-of-function phenotypes in axons

To identify factors relevant for axonal MT polymerisation, we performed a detailed loss-of-function study with a selection of candidate factors suggested by the literature (Voelzmann *et al*., 2016). We included the MT plus-end-associating factors Eb1, Shot, CLASP/Chb (Chromosome bows) and the XMAP215 homologue Msps. As MT shaft-binding candidates, we chose Tau and Map1b/Futsch. To explore the impact of tubulin availability on polymerisation, we used α1-tubulin/αtub84B (the predominant α-tubulin expressed in the fly nervous system; FlyAtlas 2, University of Glasgow, UK; A.V., unpublished) and Stathmin (a promoter of tubulin pools; (Duncan *et al*., 2013; Manna *et al*, 2007); J.F. and A.V., unpublished).

We analysed primary neurons derived from animals carrying loss-of-function mutations of the respective genes. To exclude that phenotypes are masked by maternal contribution (which is wild-type gene product deposited in the eggs by the heterozygous mothers), we used two strategies (Prokop *et al*., 2012; Sánchez-Soriano *et al*., 2010): where possible, we analysed late larval brain-derived primary neurons at 18 HIV (from now on referred to as larval neurons); if mutants did not reach larval stages, we used embryo-derived neurons that were kept for 5-7 days in pre-culture to deplete maternal product and then plated and grown for 12 hrs (pre-cultured neurons). In all cases, primary neurons were immuno-stained either for endogenous tubulin to assess axon length, or for endogenous Eb1 protein to gain a first insight into the polymerisation state of axonal MTs.

Except for Shot and Futsch, loss of all other factors displayed a significant reduction in the number of Eb1-positive plus-end comets (Figs. 1A-D, I and 1-S1A). In addition, we measured the mean intensities and mean lengths of Eb1 comets and used multiplication of these two parameters to approximate Eb1 amounts at MT plus-ends. The strong hypomorphic *Eb1^04524^* mutant allele is known to display severe, but not complete reduction of protein levels (Elliott *et al*., 2005); accordingly, Eb1 amounts at MT plus-ends were severely, but not completely diminished in neurons mutant for this allele (Figs. 1D,J and 1-S1B). Out of the remaining seven candidate factors, only two further genes showed the same qualitative mutant phenotype: *msps^A/A^* showed a reduction almost as strong as *Eb1^04524/04524^*, and *tau^KO/KO^* mutant neurons showed a milder but reliable Eb1 depletion (Fig. 1B,C, J). In all cases, the drop in Eb1 amounts was to almost equal parts due to reductions in comet length and intensity (Fig. 1-S2A,B); across all genotypes tested. They correlated well with reduced comet velocities and lifetimes when assessed in live imaging experiments (Fig. 1K,L and 1-S2C-E; using Shot-Ctail::GFP as readout for plus-end dynamics; Alves-Silva *et al*., 2012). Our data demonstrate therefore that comparable comet length/velocity correlations made *in vitro* (Roostalu *et al*, 2020) are relevant in the cellular context of axons.

**Fig. 1.**
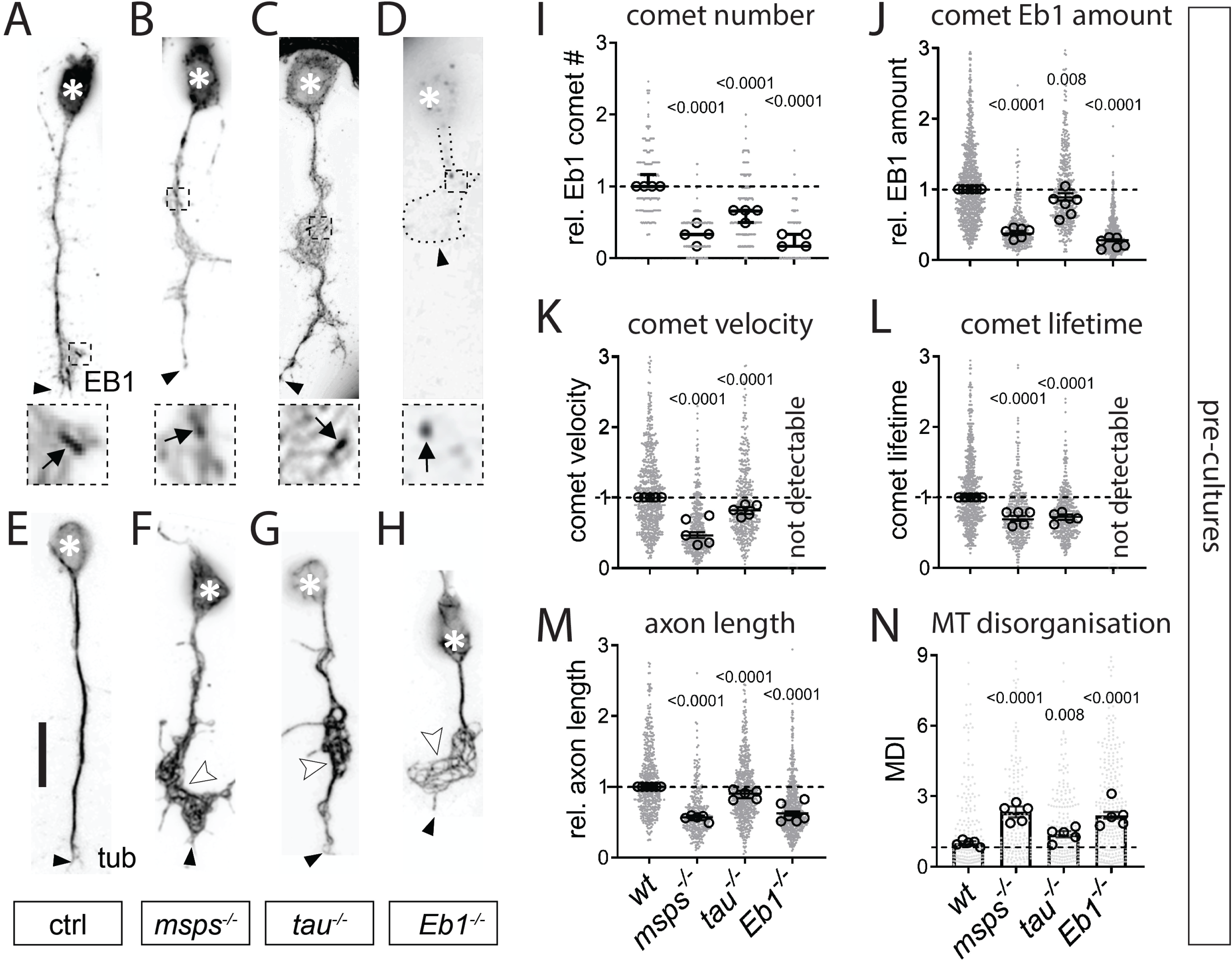
Eb1, Msps and Tau share the same combination of axonal loss-of-function phenotypes in *Drosophila* primary neurons. **A-H**) Images of representative examples of embryonic primary neurons pre-cultured to 6 days (to deplete maternal gene product; see methods) and either immuno-stained for Eb1 (top) or for tubulin (bottom); neurons were either wild-type controls (ctrl) or carried the mutant alleles *msps^1^*, *tau^KO^* or *Eb1^04524^* in homozygosis (from left to right); asterisks indicate cell bodies, black arrow heads the axon tips, white arrow heads point at areas of MT curling, dashed squares in A-D are shown as 3.5-fold magnified close-ups below each image with black arrows pointing at Eb1 comets; the axonal outline in D is indicated by a dotted line; scale bar in A represents 15 µm in all images. **I-N)** Quantification of different parameters (as indicated above each graph) obtained from pre-cultured embryonic primary neurons with the same genotypes as shown in A-H. Data were normalised to parallel controls (dashed horizontal line) and are shown as median ± 95% confidence interval (I-M) or mean ± SEM (N); data points in each plot, taken from at least two experimental repeats consisting of 3 replicates each; large open circles in graphs indicate median/mean of independent biological repeats. P-values obtained with Kruskall-Wallis ANOVA test for the different genotypes are indicated in each graph. For raw data see Tab. T1.

To assess whether reduced comet velocities or numbers correlate with impaired axon growth, we performed tubulin staining and measured axon length. Out of the eight factors, all but Futsch/MAP1B showed a decrease in axon length ranging between 10 and 43% when compared to parallel control cultures with wild-type neurons (Figs. 1E-H, M and 1-S1C). In these experiments, Tau-, Msps-, Shot- or Eb1-deficient axons showed areas of prominent disintegration of axonal MT bundles where MTs displayed intertwined, criss-crossing curls which can be quantified via the MT disorganisation index (MDI; see methods; white arrowheads in 1F-H,N and 1-S1D). This MT curling phenotype was known for Shot and Eb1 (Alves-Silva *et al*., 2012), but unexpected for Tau and Msps.

Taken together, out of eight candidate factors, loss of Futsch/MAP1B was the only condition showing no obvious defects. In contrast, loss of Eb1, Msps and Tau stood out by showing the same phenotypic pattern: reductions in comet numbers, comet velocities, Eb1 amounts and axon lengths, as well as a MT curling phenotype shared with loss of Shot. These five phenotypes appeared consistently milder in *tau* than *Eb1* or *msps* mutant neurons. Our observations raised the question as to why these three factors display such a striking pattern of collective phenotypes.

### Eb1, Msps and Tau genetically interact during axonal MT regulation both in culture and *in vivo*

To investigate potential functional relationship between Eb1, Msps and Tau, we performed genetic interaction studies. We used heterozygous conditions (i.e. one mutant and one normal copy) of genes to reduce their respective protein levels. If reduced protein levels of different genes combined in the same neurons cause a phenotype, this suggests that they are likely to function in a common pathway. For our studies, we used larval neurons and five different readouts: axon length (Fig. 2A), MT curling (Fig. 2A’), Eb1 amounts at MT plus-ends (Fig. 2A’’), comet numbers (Fig. 2-S1A) and comet dynamics (Fig. 2-S1B).

**Fig. 2.**
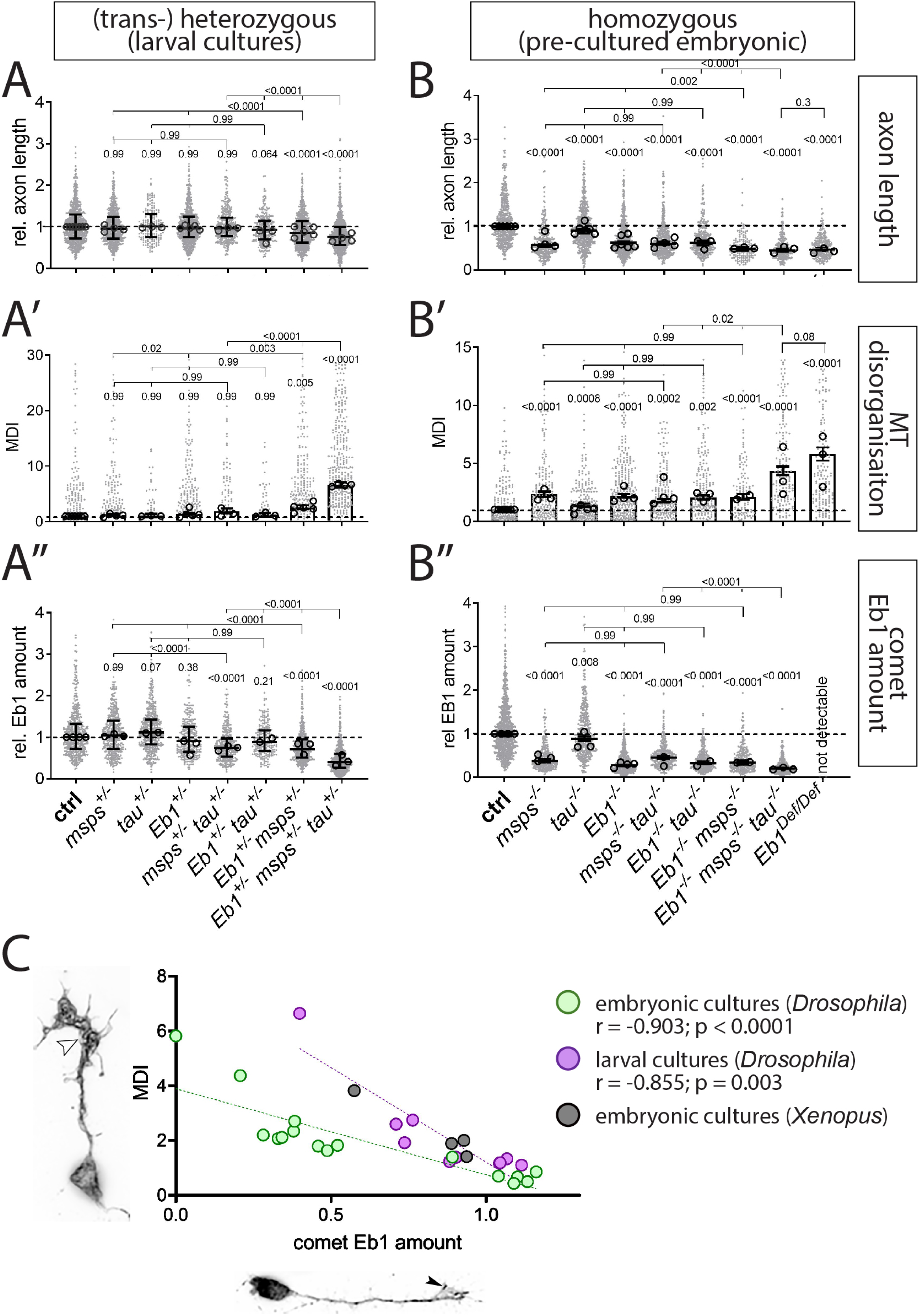
*Eb1, tau* and *msps* interact genetically. **A-B’’**) Axon length, MT curling and Eb1 amount (as indicated on the right), for primary neurons displaying heterozygous (A-A’’, larval cultures) and homozygous (B-B’, embryonic 6d pre-cultures’) mutant conditions, alone or in combination. Data were normalised to parallel controls (dashed horizontal lines) and are shown as scatter dot plots with median ± 95% confidence interval (A, A’’,B, B’’) or bar chart with mean ± SEM (A’, B’) of at least two independent repeats with 3 replicates each; large open circles in graphs indicate median/mean of independent biological repeats. P-values above data points/bars were obtained with Kruskall-Wallis ANOVA tests; used alleles: *msps^A^*, *tau^KO^*, *Eb1^04524^*. **C**) The graph compares Eb1 amounts at MT plus-ends with MT disorganisation (MDI) for a range of genetic conditions used in this work: green dots show data from pre-cultured embryonic neurons (B’ vs. B’’), purple dots show comparable data obtained from larval primary neurons (A’ vs. A’’); in addition, green/purple dots contain data from Figs. 1J and 1-S1B,D; black dots show similar data obtained from primary *Xenopus* neurons (Fig.G,H); r and p-value determined via non-parametric Spearman correlation analysis; see further detail of these correlations in Fig.2-S2. For raw data see Tab. T2.

To assess the baseline, we analysed single-heterozygous mutant neurons (*Eb1^04524/+^, msps^A/+^* or *tau^KO/+^*), none of which displayed any phenotypes. Certain double-combinations of these heterozygous conditions in the same neurons (*Eb1^04524/+^ msps^A/+^* and *tau^KO/+^ msps^A/+^*) displayed a mild but significant reduction in Eb1 amounts at MT plus-ends and a trend towards stronger MT curling. Strong enhancement of the phenotypes for all five readouts was observed in *Eb1^04524/+^ msps^A/+^ tau^KO/+^* triple-heterozygous neurons, suggesting that the three factors are functionally linked when regulating MT plus-ends, axon growth and MT organisation (Fig. 2A’-A’’ and 2-S1A,B).

The triple-heterozygous condition had a similar effect in fly brains *in vivo* when using axons of T1 medulla neurons in the adult optic lobe as readouts. In control flies analysed at 26-27 days after eclosure, axons of T1 neurons contained prominent MT bundles labelled by α-tubulin::GFP (Fig.3A; Qu *et al*., 2019). In contrast, T1 axons of triple-heterozygous mutant flies aged in parallel, showed a strong increase in areas where MTs become unbundled and twisted and axons displayed prominent swellings often containing curled MTs (Fig.3A-D). These data strongly suggest that the co-operating network of Eb1, Msps and Tau is relevant *in vivo*.

**Fig. 3.**
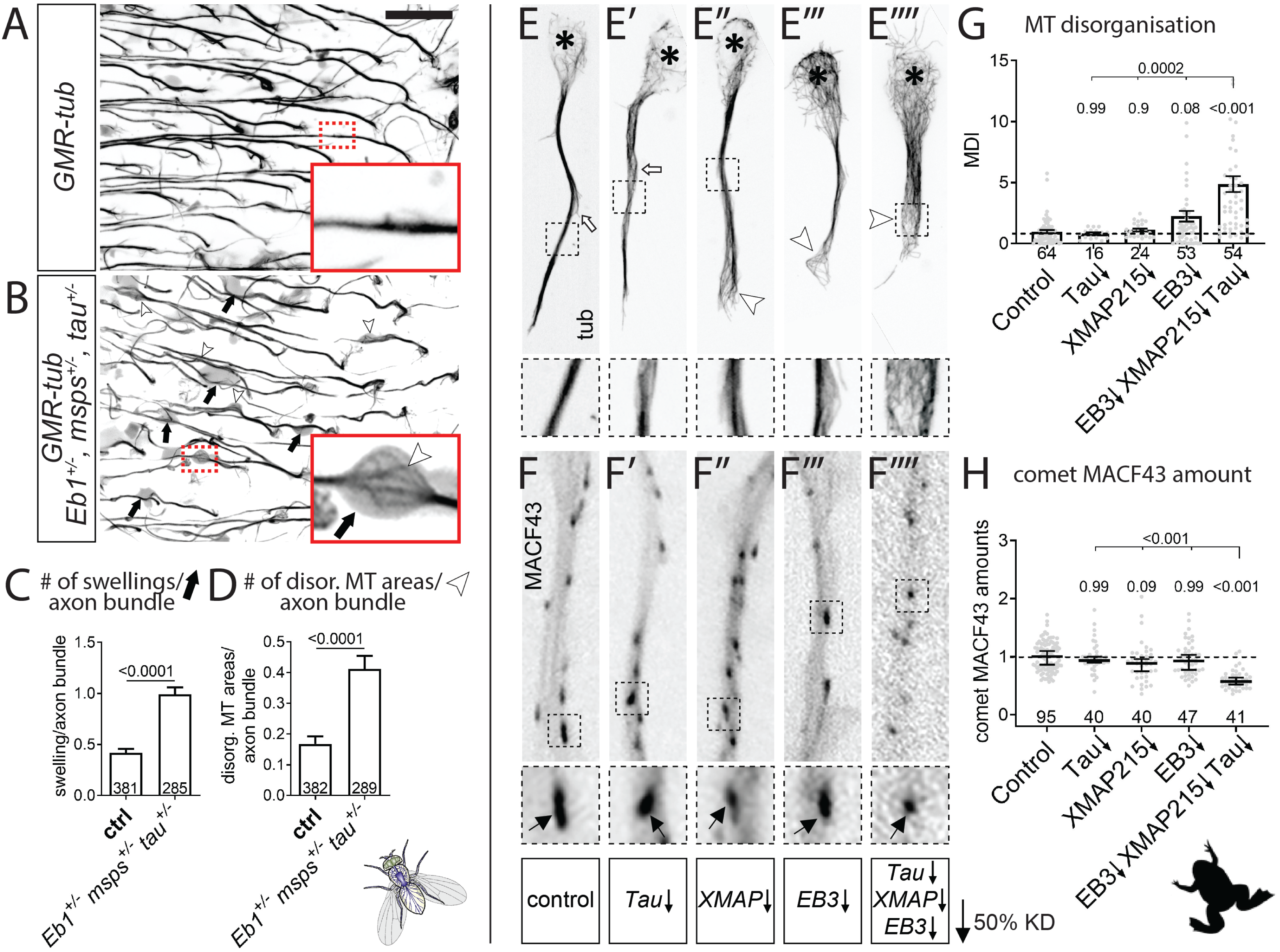
Eb1, Msps and Tau functionally interact in the fly brain and in frog primary neurons. **A,B**) Medulla region of adult brains at 26-27 days after eclosure, all carrying the *GMR31F10-Gal4* driver and *UAS-GFP-α-tubulin84B* (*GMR-tub*) which together stain MTs in a subset of lamina neuron axons that terminate in the medulla; the further genetic background is either wild-type (A) or triple-heterozygous (*Eb1^04524/+^msps^A/+^ tau^KO/KO^*; B); white/black arrows indicate axonal swellings without/with MT curling; rectangles outlined by red dashed lines are shown as 2.5 fold magnified insets where white arrow heads point at disorganised curling MTs. **C,D**) Quantitative analyses of specimens shown in A and B: relative number of total swellings per axon (C) and of swellings with MT curling per axon (D); bars show mean ± SEM; P values from Kruskal– Wallis one-way tests are given above each column, merged sample numbers (i.e. individual axon bundles) from at least two experimental repeats at the bottom of each bar. **E-E’’’’**) Primary *Xenopus* neurons stained for tubulin (tub): asterisks indicate cell bodies, white arrows indicate unbundled MTs, white arrowheads unbundled areas with MT curling. **F-F’’’**) *Xenopus* neurons labelled with MACF43::GFP (MACF43): black arrows point at comets (visible as black dots). In E-F’’’, black-stippled squares in overview images are shown as 2.5 fold magnified close-ups below; ↓ behind gene symbols indicates 50% knock-down thus approximating heterozygous conditions. **G,H**) Quantification of specimens shown in E-F’’’’ with respect to MT curling (G) and comet amount of MACF43::GFP (H); data were normalised to parallel controls (dashed horizontal lines) and are shown as mean ± SEM (G) or median ± 95% confidence interval (H); merged sample numbers from at least two experimental repeats are shown at the bottom, P-values obtained with Kruskall-Wallis ANOVA tests above data points/bars. The scale bar in A represents 15 µm in A,B and 20 µm in E-F’’’’. For raw data see Tab. T3.

### Eb1, Msps and Tau interaction is evolutionarily conserved in *Xenopus* neurons

To assess potential evolutionary conservation, we evaluated whether Eb1, Msps and Tau interact also in frog primary neurons. In the frog *Xenopus laevis*, *mapt* (*microtubule associated protein tau*) is the only *tau* homologue, *XMAP215* (*ckap5/cytoskeleton associated protein 5*) the only *msps* homologue, and *EB3* (*mapre3/microtubule associated protein RP/EB family member 3*) is the only one of three *Eb1* homologues that is prominently expressed in the nervous system (Bowes *et al*, 2009; Karimi *et al*, 2017). We used morpholinos against these three genes. Similar to our strategy in *Drosophila*, we analysed MTs by staining for endogenous tubulin (Fig.3E-E’’’), and measured Eb3 comet amounts in live movies using the Eb protein-binding peptide MACF43-Ctail::GFP as readout (comparable to Shot-Ctail::GFP used in Fig.1K,L; see Methods; Honnappa *et al*., 2009; Slater *et al*., 2019).

To approximate heterozygous mutant conditions used in our *Drosophila* experiments, we adjusted morpholino concentrations to levels that achieved knock-down of each of the three genes to ∼50% (Fig. 3-S1A-B and Lowery *et al*., 2013; Slater *et al*., 2019). Individual knock-downs to approximately 50% did not cause prominent decreases in MACF43::GFP comet amounts or increases in MT curling; but when knock-down of all three factors was combined in the same neurons, we found a reduction in MACF43::GFP comet amounts to 60% and a 4.8 fold increase in MT bundle disintegration and curling (Figs. 3E’’’,F’’’,G,H and 3-S1C,D).

Together, the three genes appear to functionally interact also in *Xenopus*, suggesting evolutionary conservation of the underpinning mechanisms.

### Eb1, Msps and Tau form an interdependent functional network with key roles played by Eb1

To investigate the underlying mechanisms, we next asked whether the functions of the three factors are hierarchically and/or interdependently organised, or whether they regulate the assessed MT properties through independent parallel mechanisms. To distinguish between these possibilities, we combined homozygous mutant conditions of the three mutant alleles in the same neurons (*Eb1^04524/04524^ msps^A/A^ tau^KO/KO^*) and asked whether this condition enhances phenotypes over single-homozygous conditions (suggesting parallel mechanisms), or whether they show no further increases (suggesting a hierarchical organisation) (Avery & Wasserman, 1992).

Since the homozygous mutant conditions do not all survive into the late larval stage, we used embryonic pre-cultured neurons for these experiments. As reference we used *Eb1^04524/04524^*, *msps^A/A^* and *tau^KO/KO^* single mutant neurons; since *Eb1^04524^* is a strong but not a total loss-of-function allele (Elliott *et al*., 2005), we added neurons homozygous for *Df(2R)Exel6050* (from now referred to as *Eb1^Df^*), a deficiency uncovering the entire *Eb1* locus (see Methods). The order of phenotypic strengths in these pre-cultured homozygous mutant neurons consistently was *Eb1^Df/Df^* > *Eb1^04524/04524^* ≥ *msps^1/1^* > *tau^KO/KO^* for all parameters assessed (Figs.2, 2-S1; see Discussion). The strong phenotypes of the *Eb1^Df/Df^* allele provided therefore a new reference bar, displaying values that were not excelled or even reached by any combination of homozygous mutants, including the triple-homozygous mutant neurons (Fig. 2B-B’’).

These results are consistent with a model where the three proteins co-operate interdependently, through a modality in which Tau has the weakest contribution and Eb1 is the most essential factor. This is also suggested when extracting data from a range of different genetic conditions analysed throughout this study and plotting their Eb1 amounts at MT plus-ends against the degree of MT curling in each condition (Fig.2-S2A). This plot revealed a highly significant inverse correlation between Eb1 comet amounts and MT disorganisation (Figs. 2C and 2-S2B), suggesting that Eb1 is the key factor within the functional trio linking out to MT bundle promotion.

### Eb1 and Msps depend on each other for MT plus-end localisation

We next investigated the mechanistic links between the three proteins, starting with Msps and Eb1. When co-expressing Msps::GFP together with Eb1::RFP in *Drosophila* embryonic primary neurons, both proteins prominently localised at the same MT plus-ends, with Msps::GFP localising slightly distal to Eb1 (Fig. 4A-A’’, Movie M1). Their localisation at the same MT plus-ends was even clearer in kymographs of live movies where both proteins remained closely associated during comet dynamics (Fig.4B,B’). These findings agree with localisation data for Eb proteins and XMAP215 *in vitro* (Maurer *et al*, 2014), suggesting that the same mechanisms apply in the cellular context of axons and might explain the linked phenotypes we observed for the two factors. This was further supported by their co-dependence for achieving prominent plus-end localisation:

**Fig. 4.**
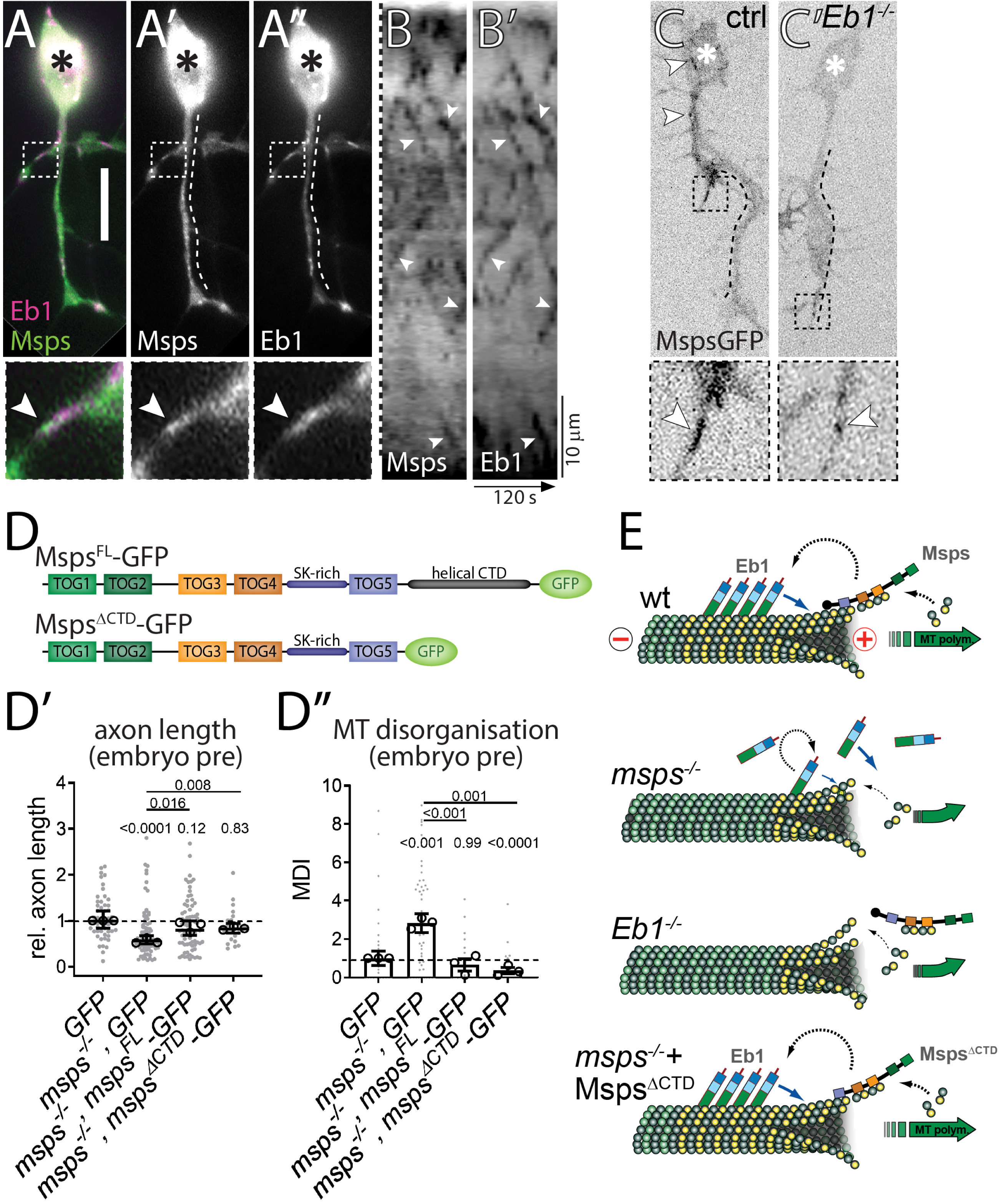
Eb1 and Msps depend on each other for MT plus-end localisation. **A-A’’**) Primary neurons at 6HIV co-expressing Eb1::mCherry (magenta, Eb1) and Msps^FL^::GFP (green, Msps) and imaged live; asterisks indicate somata, scale bar represents 10µm, dashed boxes indicate the positions of the 3.5-fold magnified close-ups shown at the bottom with arrowheads pointing at the position of Msps::GFP accumulation (same in C,C’). **B,B’**) Kymograph of live movies (as in A-A’’) with the dashed line on the left representing a straightened version of the dashed lines shown in A’ and A’’ (i.e. the length of the axon; proximal at the top) and the x-axis indicating time; arrowheads point at trajectories of Msps and Eb1 which are almost identical. **C**) Primary neurons expressing Msps::GFP and imaged live, either displaying wild-type background (ctrl) or being homozygous mutant for *Eb1^04524^* (*Eb1^-/-^*); white arrowheads point at Msps::GFP comets which are much smaller in the mutant neurons. **D**) Schematic representations of Msps^FL^::GFP and Msps^ΔCTD^::GFP. **D’,D’’**) Graphs displaying axon length and MT curling (as indicated) for pre-cultured embryonic primary neurons expressing GFP or Msps::GFP constructs via the *elav-Gal4* driver, either in wild-type or *msps^A/1^* mutant background; data were normalised to parallel controls (dashed horizontal lines) and are shown as median ± 95% confidence interval (D’) or mean ± SEM (D’’) from at least two experimental repeats; large open circles in graphs indicate median/mean of independent biological repeats. P-values obtained with Kruskall-Wallis ANOVA tests are shown above data points/bars. **E**) Model view of the results shown here and in Fig. 1; note that yellow circles represent GTP-tubulin which mediates the binding of Eb1; for further explanations see main text and Discussion. For raw data see Tab. T4.

We already reported that loss of Msps causes severe Eb1 depletion and comet shortening at MT plus-ends (Fig1. 1B, D and 1-S1B). This is likely due to the role of Msps as a tubulin polymerase (Brouhard *et al*., 2008) which helps to sustain a prominent GTP-tubulin cap to which EB1 can bind (Zanic *et al*., 2013). To test this, we stained *msps^1/1^* mutant neurons for GTP-tubulin, and found that GTP caps displayed reduced lengths comparable to the shortened Eb1 comets (Figs. 1J *vs*. 5-S2C). This result, together with our finding of reduced comet velocity in *msps^1/1^* mutant neurons (Fig. 1K), is consistent with the hypothesis that Msps-dependent polymerisation increases Eb1 amounts at MT plus-ends (see Discussion).

*Vice versa*, we expressed Msps::GFP in *Eb1^04524/04524^* mutant neurons, which likewise revealed severe depletion of Msps at growing MT plus-ends when compared to controls (Fig. 4C,C’; Movies M2 and M3). These results suggest that Msps/XMAP215 can bind MT plus-ends independently, but that its binding is enhanced if Eb1 is present (see also Maurer *et al*., 2014; Zanic *et al*., 2013). One mechanism through which Eb1 might recruit Msps/XMAP215 at MT plus-ends is through adaptors such as SLAIN in vertebrates or TACC (Transforming acidic coiled-coil protein) or Sentin in non-neuronal *Drosophila* cells (Brouhard *et al*., 2008; Lee *et al*, 2001; Li *et al*, 2011; Li *et al*., 2012; Lowery *et al*., 2013; Tang *et al*, 2020; van der Vaart *et al*, 2012). However, our functional studies using homozygous mutant neurons carrying *tacc^1^* and *sentin^ΔB^* loss-of-function mutant alleles failed to display axon shortening or MT curling, thus arguing against a prominent role of these factors during Eb1-dependent Msps recruitment (details in Fig.4-S1A,B). To further substantiate these findings, we generated a *msps^ΔCterm^-GFP* construct which lacks the C-terminal domain essential for the interaction with adaptors (Fig.4D; Fox *et al*, 2014; Mortuza *et al*, 2014). When *msps^ΔCterm^-GFP* or *msps^FL^-GFP* full length controls were transfected into *msps^A/146^* mutant neurons, we found that both protein variants were similarly able to improve axon length and MT curling defects (Fig.4D,D’), thus likewise arguing against the requirement of adaptors to recruit Msps to MT plus-ends in fly neurons.

In conclusion, our data clearly demonstrate that Eb1 and Msps require each other to achieve prominent MT plus-end localisation in axons: Msps likely maintains Eb1 at MT plus-ends through promoting GTP-cap formation, as is supported by our findings with GTP-tubulin staining. *Vice versa,* Eb1-mediated recruitment of Msps might involve roles of Eb1 in the structural maturation of MT plus-ends (see Discussion; Maurer *et al*., 2014; Zanic *et al*., 2013).

### Tau promotes Eb1 pools at MT plus-ends by outcompeting it from lattice binding

Similar to Msps deficiency, we found that also loss of Tau leads to a reduction of Eb1 comet sizes in both *Drosophila* (Fig.1C,J and 1-S2A,B) and *Xenopus* neurons (Fig.3-S1G). In wild-type fly neurons, Tau localises along MT lattices and appears not to extend into the Eb1 comet at the MT plus-end (Fig.5A’A’’). This distribution is consistent with reports that Tau and Eb1 do not overlap at MT plus-ends due to their respective higher affinities for GDP- and GTP-tubulin (Castle *et al*, 2020; Duan *et al*, 2017), suggesting that Tau promotes Eb1 plus-end localisation through indirect mechanisms not involving their physical interaction.

**Fig. 5.**
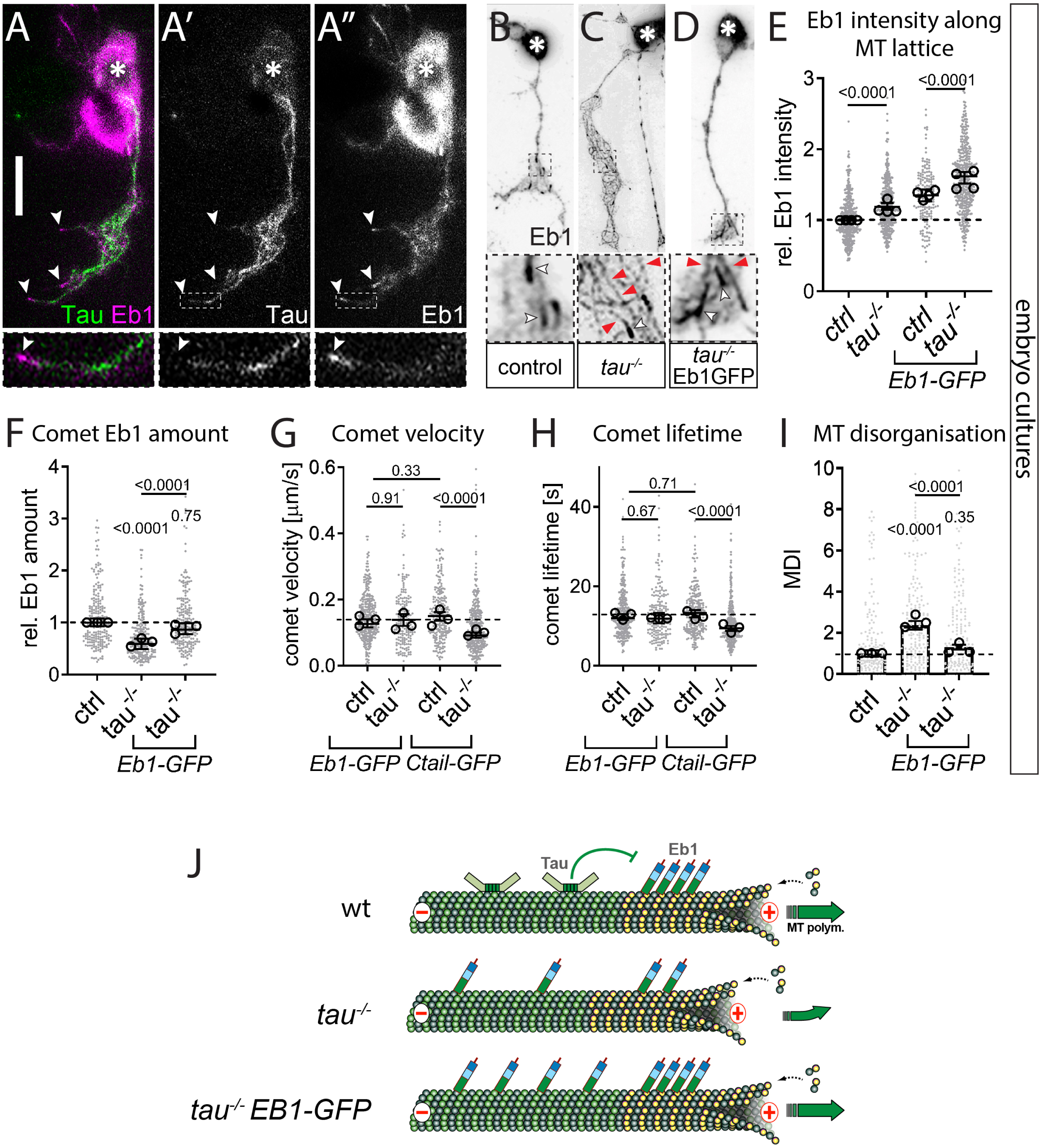
Tau promotes Eb1 pools at MT plus-ends by outcompeting Eb1’s association with the MT lattice. **A-A’’**) Example of an embryonic neuron at 6 HIV imaged live with curling MTs to illustrate Tau binding (green) along the MT lattice, separated from Eb1 comets (magenta); asterisks indicate the soma, the scale bar represents 10µm, dashed boxes indicate the positions of the 4-fold magnified close-ups shown at the bottom, white arrowheads point at Eb1 comets (same in B-D). **B-D**) Primary neurons at 6 HIV stained for Eb1 which are either wild-type (B) or mutant for *tau^KO/Df^* (C) or expressing Eb1::GFP driven by *elav-Gal4* in *tau^KO/Df^* mutant background (D); white arrowheads indicate Eb1 comets, red arrowheads Eb1 lattice localisation. **E-I**) Different parameters (as indicated) of control (ctrl) or *tau^KO/Df^* (*tau^-/-^*) mutant neurons at 6 HIV without/with *elav-Gal4*-driven expression of Eb1::GFP or Shot-Ctail::GFP (as indicated); data were normalised to parallel controls (dashed horizontal lines) and are shown as scatter dot plots with mean ± SEM (I) or median ± 95% confidence interval (E-H) from at least two independent repeats with 3 experimental replicates; large open circles in graphs indicate median/mean of independent biological repeats. P-values obtained with Kruskal-Wallis ANOVA and Dunn’s posthoc test as shown above data points/bars. **J**) Model view of the results shown here; note that yellow circles represent GTP-tubulin which provides higher affinity for Eb1 binding; for further explanations see main text and Discussion. For raw data see Tab. T5.

We noticed that the reduction of Eb1 comet sizes in Tau-deficient neurons is accompanied by a 20% increase in the intensity of Eb1 along MT lattices (Fig.5C,E for 6 day pre-culture, Fig.5-S2A for embryonic cultures). This effect is specific for *tau* and not observed in *msps^A/A^* mutant neurons (Fig. 5-S2A,B). Tau has previously been shown to protect MTs against Katanin-induced damage (Qiang *et al*, 2006). Increased MT repair upon loss of Tau could therefore promote the incorporation of GTP-tubulin along the lattice which would, in turn, recruit Eb1 (Vemu *et al*, 2018). However, MT lattices in Tau-deficient neurons did not show obvious increases in GTP-tubulin (Fig. 5-S1B, D), arguing against the repair hypothesis. In the same specimens, GTP-tubulin amounts at MT plus-ends were clearly reduced, consistent with the observed Eb1 comet depletion in Tau-deficient neurons (Figs. 1J *vs*. 5-S1A-C).

We observed that over-expression of Tau in wild-type neurons decreased Eb1 levels along MT lattices even further (Fig. 5-S1E), and similar observations were reported *in vitro* with mammalian versions of the two proteins (Ramirez-Rios *et al*, 2016). We hypothesised therefore that Tau may competitively prevent lattice-binding of Eb1, as similarly observed for other MAPs (Qiang *et al*, 2018): if Tau is absent, MT lattices would therefore turn into a sink for Eb1 and sequester Eb1 pools away from MT plus-ends. Such a mechanism could be particularly relevant in narrow axons which display a high relative density of MTs (Prokop, 2020). In support of our hypothesis, we found that *tau^KO/KO^* mutant neurons overexpressing Eb1::GFP showed even greater Eb1 intensity along MT lattices, but also replenished Eb1 amounts at MT plus-ends (Fig. 5D-F). This treatment was sufficient to suppress Tau-deficient phenotypes, as reflected in the recovery of Eb1 comet dynamics (improved velocity and lifetime) and strongly reduced MT curling (’Eb1-GFP’ in Fig. 5G-I).

These effects were specific for Tau and Eb1: Firstly, no rescue of *tau^KO/KO^* mutant phenotypes was observed when expressing Shot-Ctail::GFP (as used in Fig.1K,L) to track MT plus-ends without altering Eb1 levels (’Ctail-GFP’ in Fig.5G,H). Secondly, Eb1::GFP could not restore MT plus-end dynamics in *msps^A/A^* mutant neurons (Fig. 5-S2D,E), a finding that provides important insights also into the hierarchical relationships between Eb1, Msps and Tau (see Discussion).

In conclusion, we propose that Tau contributes to MT polymerisation dynamics and MT organisation indirectly, through preventing that Eb1 is sequestered away from MT plus-ends. The fact that Eb1::GFP can rescue *tau* mutant phenotypes including their MT curling, further highlights Eb1’s crucial role in MT bundle promotion (compare Fig.2C, Fig.2-S2) and may suggest it even as a potential therapeutic target (see Discussion).

### An Eb1- and spectraplakin-dependent guidance mechanism explains roles of the functional trio in MT bundle organisation

Since Eb1 amounts at MT plus-ends inversely correlate with MT curling (Figs.2C and 2-S2), bundle deterioration in either Eb1-, Msps- or Tau-deficient neurons might be a consequence of their negative impacts on Eb1 localisation. Eb1 at polymerising MT plus-ends has been proposed to recruit the C-terminus of Shot which simultaneously binds cortical F-actin via its N-terminus (Fig.6A). Through this cross-linking activity, Shot has been proposed to guide the extension of MTs along the axonal surface into parallel bundles (Fig.6F); accordingly, depletion of either Shot or Eb1 causes MT curling; Alves-Silva *et al*., 2012; Sánchez-Soriano *et al*., 2009; Voelzmann *et al*, 2017); Figs.1D, 6E).

**Fig. 6.**
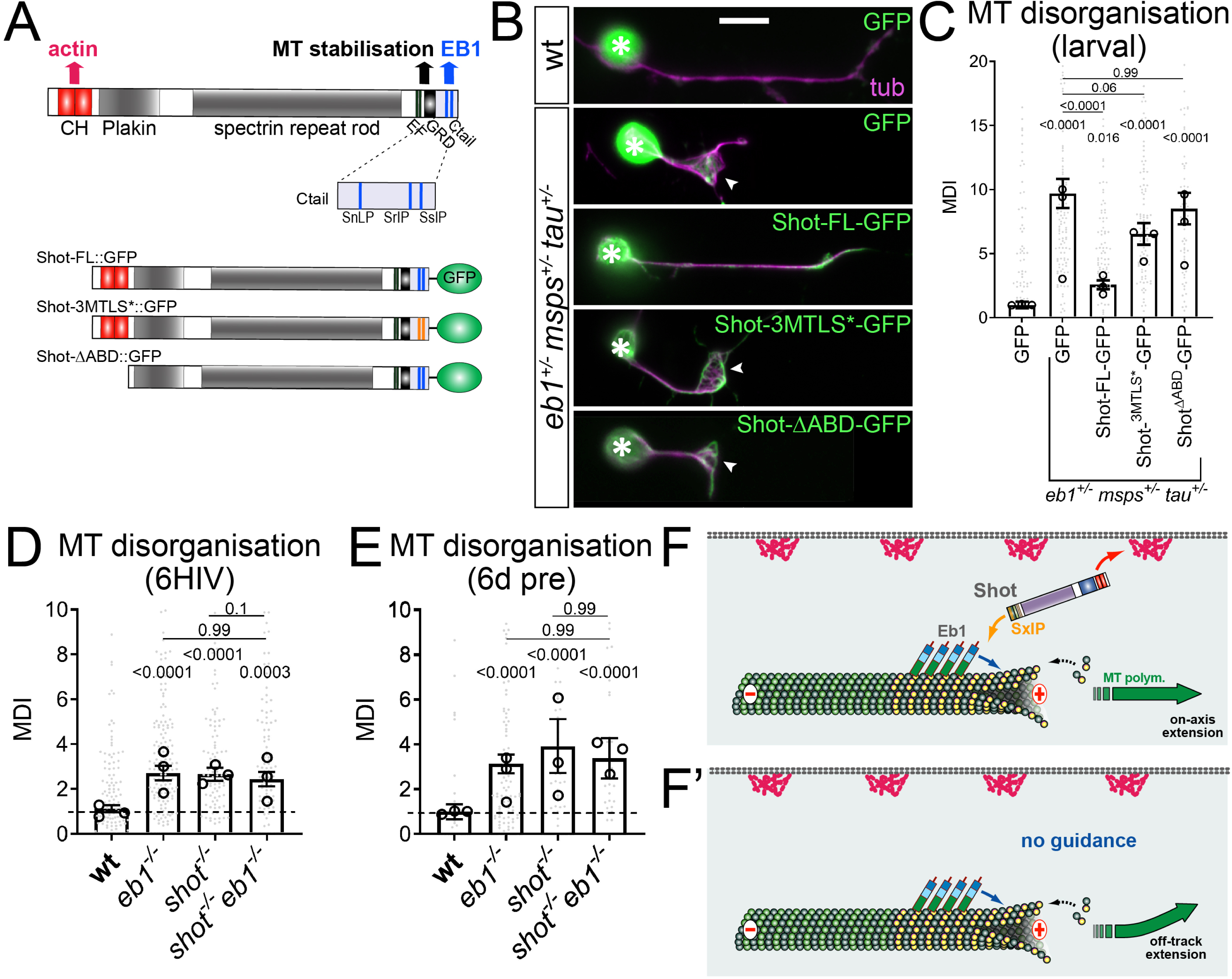
Shot-mediated guidance mechanistically links Eb1 at MT plus-ends to bundle organisation. **A**) Schematic representation of Shot constructs (CH, actin-binding calponin-homology domains; EF, EF-hand motifs; GRD, MT-binding Gas2-related domain; Ctail, unstructured MT-binding domain containing Eb1-binding SxIP motifs in blue); in Shot-3MtLS*::GFP the SxIP motifs are mutated. **B**) Fixed primary neurons at 18HIV obtained from late larval CNSs stained for GFP (green) and tubulin (magenta), which are either wild-type (top) or *Eb1^04524/+^msps^A/+^ tau^KO/+^* triple-heterozygous (indicated on right) and express GFP or either of the constructs shown in D; scale bar 10µm. **C**) Quantification of MT curling of neurons as shown in B. **D,E**) MT curling in *shot^3/3^ Eb1^04524/04524^* double-mutant neurons is not enhanced over single mutant conditions assessed in fixed embryonic primary neurons at 6HIV (D) or 12HIV following 6 day pre-culture (E). In all graphs data were normalised to parallel controls (dashed horizontal lines) and are shown as mean ± SEM from at least two independent repeats with 3 experimental replicates each; large open circles in graphs indicate median/mean of independent biological repeats. P-values obtained with Kruskall-Wallis ANOVA tests are shown above bars. **F,F’**) Model derived from previous work (Alves-Silva *et al*., 2012), proposing that the spectraplakin Shot cross-links Eb1 at MT plus-ends with cortical F-actin, thus guiding MT extension in parallel to the axonal surface; yellow dots represent GTP-tubulins providing high affinity sites for Eb1-binding.

In support of this guidance hypothesis, we found that severe MT curling observed in *Eb1^+/-^ msps^+/-^ tau^+/-^* triple-heterozygous mutant neurons (which have strongly reduced Eb1 comet amounts; Fig.2A’’) was significantly improved from 7.3-fold (with GFP-expression) to 1.4-fold, when over-expressing full length Shot-FL::GFP (Fig.6B-C; both compared to wild-type controls). This rescue could also be mediated through Eb1-independent roles of Shot in MT stabilisation mediated by the C-terminal Gas2-related domain (Fig.6A; Alves-Silva *et al*., 2012). We therefore repeated the rescue experiments with two different Shot variants that maintain MT-stabilising activity whilst specifically abolishing actin-Eb1 cross-linkage (Fig.6A,F): (1) Shot^ΔABD^::GFP lacks the N-terminal calponin homology domain required for interaction with the actin cortex; (2) Shot^3MtLS*^::GFP carries mutations in the C-terminal SxIP motifs required for Eb1 binding. Both these Shot variants failed to rescue MT curling in *Eb1^04524/+^ msps^1/+^ tau^KO/+^* triple-heterozygous mutant neurons (Fig.6B,C), hence lending further support to the guidance hypothesis based on Eb1-Shot interaction. This conclusion is further substantiated by the finding that MT curling observed in *Eb1^04524/04524^* mutant neurons is not enhanced in *Eb1^04524/04524^ shot^3/3^* double-mutant neurons, neither in normal embryonic nor in embryonic pre-cultured neurons (Figs. 6D, E).

We conclude that co-operative promotion of MT polymerisation through Eb1, Msps and Tau appears to perform its additional function in axon bundle organisation through Shot-mediated MT guidance downstream of Eb1. In this way, the three proteins uphold the numbers/mass as well as the bundled organisation of MTs as two key features underpinning the formation and long-term maintenance of axons.

## Discussion

### New understanding of the role and regulation of MT polymerisation and guidance in axons

Understanding the machinery of MT polymerisation is of utmost importance in axons where MTs form loose bundles of enormous lengths which are essential for axonal morphogenesis and serve as life-sustaining transport highways (Prokop, 2020). These bundles must be maintained in functional state for up to a century in humans (Hahn *et al*., 2019). To achieve this, MT polymerisation is required to generate MTs *de novo*, repair or replace them. The underpinning machinery is expected to be complex (Voelzmann *et al*., 2016), but deciphering the involved mechanisms will pay off by delivering new strategies for tackling developmental and degenerative axon pathologies (Prokop, 2021).

Here we made important advances to this end. Having screened through 13 candidates (this work and Beaven *et al*., 2015), we found the three factors Eb1, Msps and Tau to stand out by expressing the same combination of phenotypes, and by displaying functional interaction in both *Drosophila* and *Xenopus* neurons. We found that their functions are not only important to maintain MT polymerisation, but also to align MTs into parallel arrangements, thus contributing in two ways to MT bundle formation and maintenance, both in culture and *in vivo*. The observed impact on MT organisation is also consistent with roles of XMAP215 during MT guidance in growth cones of frog neurons (Slater *et al*., 2019).

Our data reveal that various mechanisms observed *in vitro* or in non-neuronal contexts, apply in the biological context of axons, which was unpredictable for two reasons: Firstly, of the three proteins only the human tau homologue has OMIM®-listed links to human axonopathies, and these do not necessarily relate to MT polymerisation (Morris *et al*., 2011). Secondly, axons and non-neuronal cells can display significant mechanistic deviations as shown for CLIP170/190 (Beaven *et al*., 2015) and for the MT localisation of Msps which is facilitated by Sentin or dTACC in non-neuronal cells, dendrites and *in vitro* (Lee *et al*., 2001; Li *et al*., 2011; Li *et al*., 2012; Tang *et al*., 2020) but seemingly not in axons (Fig.4S1). However, other mechanisms we observed in axons matched previous reports: (1) the complementary binding preferences of Eb1 and Tau for GTP-/GDP-tubulin (Castle *et al*., 2020; Zanic *et al*, 2009); (2) the mutual enhancement of Eb1 and XMAP215/Msps (Kronja *et al*, 2009; Li *et al*., 2012; Zanic *et al*., 2013); (3) the correlation of GTP cap size with comet velocity (Roostalu *et al*., 2020). Furthermore, we observed that depletion of α1-tubulin in neurons mutant for *αtub84B* or *stathmin* (a promoter of tubulin availability; (Duncan *et al*., 2013; Manna *et al*., 2007); J.F. and A.V., unpublished) affects comet numbers but not comet dynamics (Fig.1-S1); this is consistent with observations that MT nucleation *in vitro* is far more sensitive to tubulin levels than polymerisation (Consolati *et al*, 2020).

### Eb1 and XMAP215/Msps are core factors promoting MT polymerisation and guidance

Apart from demonstrating the relevance of various molecular mechanisms in the context of axonal MT regulation, our work provides key insights as to how they integrate into one consistent mechanistic model of biological function (summarised in Fig.7).

**Fig. 7.**
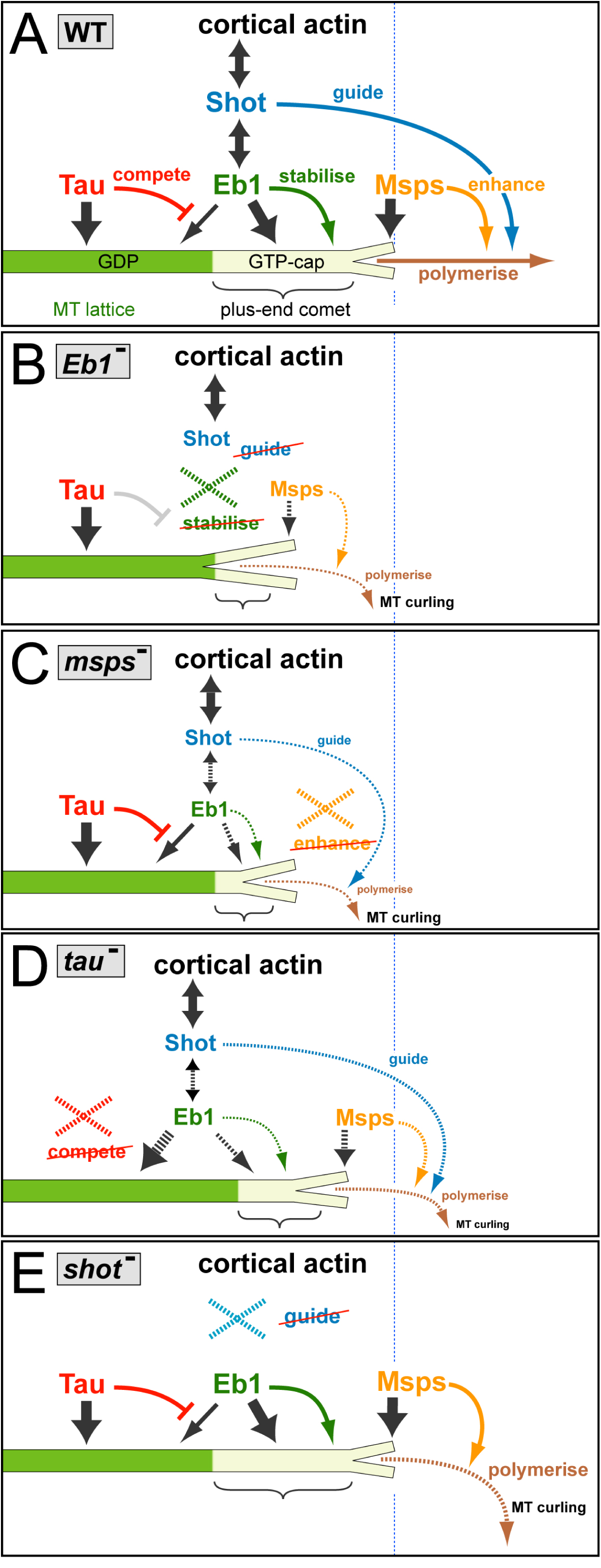
A mechanistic model consistent with all reported data. **A**) In wild-type (WT) neurons, the three factors bind (dark grey arrows) to MTs at different locations: Msps binds to the very tip of MT plus-ends, Eb1 forms plus-end comets (curly bracket) by binding with higher affinity to GTP-tubulin (GTP-cap, light beige) but lagging behind the front, and Tau localises along the MT lattice primarily composed of GDP-tubulin (green). Tau outcompetes (red T-bar) low-affinity binding of Eb1 along the lattice, thus maintaining more Eb1 at MT plus-ends. Msps enhances (orange arrow) MT polymerisation (brown), thus sustaining a prominent GTP-cap that Eb1 can bind to. Eb1 stabilises MT plus-ends (green arrow) by promoting sheet formation of protofilaments at the plus-end (short Y-shaped tip), thus improving conditions for Msps binding. Eb1 also recruits the C-terminus of Shot which binds cortical actin via its N-terminus, thus establishing cross-linkage that can guide (blue arrow) extending MT plus-ends into parallel bundles. **B-E**) Illustrations explaining the changes triggered by the loss of the different factors (stippled X); any affected processes are shown as stippled lines with reduced thickness and reduced font sizes of accompanying texts. **B**) Upon loss of Eb1, the plus-end sheet structure is weakened (larger Y shape), thus reducing Msps binding and, in turn, reducing polymerisation; Shot detaches, thus abolishing guidance. **C**) Loss of Msps abolishes enhanced polymerisation so that the GTP-cap shrinks and less Eb1 binds, thus also weakening Shot binding and MT guidance. **D**) Upon loss of Tau, more Eb1 can bind to the MT lattice, thus reducing the amounts available for plus-end association; this causes a modest Eb1 depletion phenotypes with consequences similar to B, but less pronounced. **E**) Upon loss of Shot, the localisations and functions of the other three proteins are unaffected, but MT guidance is abolished.

The TOG-domain protein XMAP215/Msps is relevant for neuronal morphogenesis in fly and *Xenopus* (Lowery *et al*., 2013; Tang *et al*., 2020), likely through its expected function as a MT polymerase (Al-Bassam & Chang, 2011; Brouhard *et al*., 2008; Fox *et al*., 2014; Howard & Hyman, 2009; Li *et al*., 2012; Zanic *et al*., 2013). In contrast, *Drosophila* and vertebrate Eb proteins are only moderate promoters of MT polymerisation *in vitro* (Li *et al*., 2012; Ramirez-Rios *et al*., 2016; Zanic *et al*., 2013) and references within), but rather act as scaffolds (Akhmanova & Steinmetz, 2015). Conserved binding partners of Eb proteins are the spectraplakins which can guide extending MT plus-ends along actin networks in axons and non-neuronal cells (Alves-Silva *et al*., 2012; Voelzmann *et al*., 2017; Wu *et al*, 2008).

We propose therefore (see Fig.7A) that Eb1 is the key mediator of MT guidance into bundles (as supported by data throughout this work; Figs.2C, 2-S2, 6), and Msps the key promoter of MT polymerisation. To execute these functions, both proteins depend on each other: MT plus-end localisation of Msps is reduced upon loss of Eb1 (Fig.7B) and *vice versa* (Fig.7C). This mutual dependency is unlikely to involve their physical interaction, since MT plus-end localisation of Eb1 is known to occur tens of nanometres behind XMAP215 (Maurer *et al*., 2014; Zanic *et al*., 2013), as seems to be the case also for axonal MTs (Fig.4A). Furthermore, our data do not support an obvious role of the Sentin or dTACC adaptors in mediating Eb1-XMAP215 interactions (Figs.4D-D’’ and 4-S1).

Potential indirect mechanisms explaining this co-dependency are provided by the promotion of MT polymerisation through XMAP215/Msps which maintains a prominent GTP-cap (Fig.5-S1) that, in turn, mediates Eb1 binding (see also (Maurer *et al*., 2014; Zanic *et al*., 2013); Fig.7C).

Restricted GTP-cap formation as a limiting factor for Eb1 binding would also explain why Eb1 over-expression fails to rescue Msps-deficient phenotypes (Fig.5-S2D-F). *Vice versa*, Eb proteins promote lateral protofilament contacts which could assist in sheet formation at the very plus tip (Fig.7B), thus facilitating the binding of XMAP215/Msps (Maurer *et al*., 2014; Zanic *et al*., 2013).

### Tau contributes through an indirect mechanism of competitive binding to MT lattices

Tau and Map1b/Futsch are known to bind along MT lattices, to promote MT polymerisation *in vitro*, and to enhance axon growth in mouse and fly neurons through mechanisms that remain unclear (Brandt & Lee, 1993; Caceres & Kosik, 1990; Cleveland *et al*, 1977; DiTella *et al*, 1996; Drechsel *et al*., 1992; Hummel *et al*., 2000; Kadavath *et al*, 2015; Kiris *et al*, 2010; Levy *et al*, 2005; Liu *et al*., 2015; Panda *et al*, 1995; Ramirez-Rios *et al*., 2016; Takei *et al*, 2000; Tymanskyj *et al*, 2012; Villarroel-Campos & Gonzalez-Billault, 2014).

In our cellular model, loss of Map1b/Futsch has no obvious effects, whereas Tau shares all assessed loss-of-function mutant phenotypes with Msps and Eb1, though with weaker expression. Of these phenotypes, reduced MT plus-end localisation of Eb proteins upon loss of Tau function (Figs.1C,J) was likewise reported for frog neurons (Fig.3-S1G-I), N1E-115 mouse neuroblastoma cells and primary mouse cortical neurons (Sayas *et al*, 2015).

One proposed mechanism involves direct interaction where Tau recruits Eb proteins at MT plus ends (Sayas *et al*., 2015), consistent with other reports that Tau can bind Eb1 (Buey *et al*, 2011; Duan *et al*., 2017). However, further reports argue against overlap of Tau and Eb1 and rather show that Eb1 and Tau have complementary preferences for GTP- and GDP-tubulin, respectively (Castle *et al*., 2020; Duan *et al*., 2017; Zanic *et al*., 2009); this is also consistent with our data (Fig.5A). Furthermore, a recruitment model is put in question by our finding that Eb1 lattice localisation increases rather than decreases upon loss of Tau (Fig.5E), as similarly observed for mammalian tau and Eb1 *in vitro* (Ramirez-Rios *et al*., 2016).

Therefore, we propose a different mechanism based on competitive binding where Tau’s preferred binding to GDP-tubulin along the lattice prevents Eb1 localisation (Fig.5), comparable to Tau’s role in preventing MAP6 from binding in certain regions of the MT lattice (Baas & Qiang, 2019; Qiang *et al*., 2018). Given the high density of MTs in the narrow space of axons, lattice binding could generate a sink large enough to reduce Eb1 levels at MT plus-ends, and our rescue experiments with Eb1::GFP overexpression strongly support this notion (Fig.5). In this way, loss of Tau generates a condition comparable to a modest Eb1 loss-of-function mutant phenotype, thus explaining why Tau shares its repertoire of loss-of-function phenotypes with Msps and Eb1, but with more modest presentation. This competition mechanism might apply in axons with high MT density, for example explain the reduction in axonal MT numbers upon Tau deficiency in *C. elegans* (Krieg *et al*, 2017); it might be less relevant in larger diameter axons of vertebrates where MT densities are low (Prokop, 2020).

### Main conclusions and future perspectives

Here we have used a standardised *Drosophila* neuron system amenable to combinatorial genetics to gain understanding of MT regulation at the cellular level in axons. We propose a consistent mechanistic model that can integrate all our data, mechanisms reported in the literature, and our previous mechanistic model explaining Eb1/Shot-mediated MT guidance (Alves-Silva *et al*., 2012; Qu *et al*., 2019). This understanding offers new opportunities to investigate the mechanisms behind other important observations.

For example, the presence of an axonal sleeve of cortical actin/spectrin networks was shown to be important to maintain MT polymerisation, likely relevant in certain axonopathies (Prokop, 2021; Qu *et al*., 2017); the underlying mechanisms are now far easier to dissect. As another example, we found that loss of either Eb1, XMAP215/Msps or Tau all caused a reduction in comet numbers, consistent with reports of nucleation-promoting roles of XMAP215 (Flor-Parra *et al*, 2018; Roostalu *et al*, 2015; Thawani *et al*, 2018; Wieczorek *et al*, 2015). This might offer opportunities to investigate how MT numbers can be regulated in reproducible, neuron/axon-specific ways, thus addressing a fundamental aspect of axon morphology (Prokop, 2020).

By gradually assembling molecular mechanisms into regulatory networks that can explain axonal MT regulation at the cellular level, i.e. the level at which diseases become manifest, our studies come closer to explaining axonal pathologies which can then form the basis for the development of remedial strategies (Hahn *et al*., 2019).

## Acknowledgements

This work was made possible through support by the BBSRC to A.P (BB/I002448/1, BB/P020151/1, BB/L000717/1, BB/M007553/1) to N.S.S. (BB/M007456/1, BB/R018960/1), by the Leverhulme Trust to I.H. (ECF-2017-247), by the German Research Council (DFG) to A.V. (VO 2071/1-1), by NIH to L.A.L (R01 MH109651), and a postdoctoral fellowship from CONICYT to P.G.S. The Manchester Bioimaging Facility microscopes used in this study were purchased with grants from the BBSRC, The Wellcome Trust and The University of Manchester Strategic Fund. The Fly Facility has been supported by funds from The University of Manchester and the Wellcome Trust (087742/Z/08/Z). We thank Hiro Ohkura for kindly providing DmEb1 antibody and unpublished mutant alleles of *msps*. Stocks obtained from the Bloomington *Drosophila* Stock Center (NIH P40OD018537) were used in this study.

---

**Fig. 1-S1.**
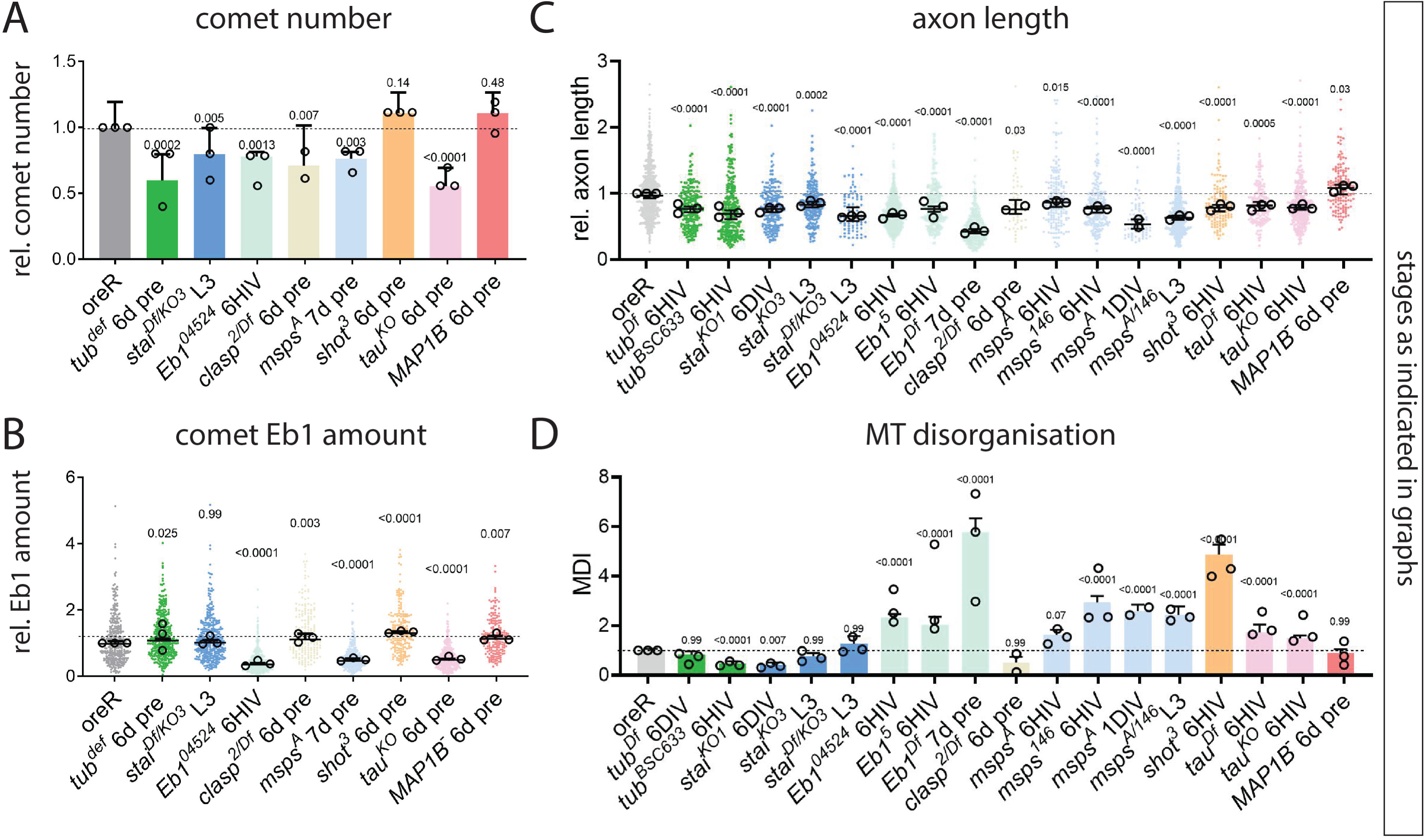
A candidate screen of axonal loss-of-function phenotypes in primary neurons. Graphs show extended data sets for four of the parameters displayed in Fig. 1 (indicated above each graph). Data points/bars representing mutant conditions for different genes are consistently colour-coded in all graphs, and conditions used are indicated below (6HIV, cultured from embryos for 6hrs; 6d/7d pre, cultured from embryos for 12hrs following 6 or 7 days pre-culture; L3, cultured from late larval CNS for 18hrs). Allele names are given as superscript: absence of slash indicates homozygous, presence of slash hetero-allelic conditions. Data were normalised to parallel controls (dashed horizontal line) and are shown as median ± 95% confidence interval (B,C) or mean ± SEM (A,D); data points from at least two experimental repeats consisting of 3 replicates each are shown, large open circles in graphs indicate median/mean of independent biological repeats. P-values obtained with Kruskall-Wallis ANOVA test above data points/bars. For raw data see Tab. T1-S1.

**Fig. 1-S2.**
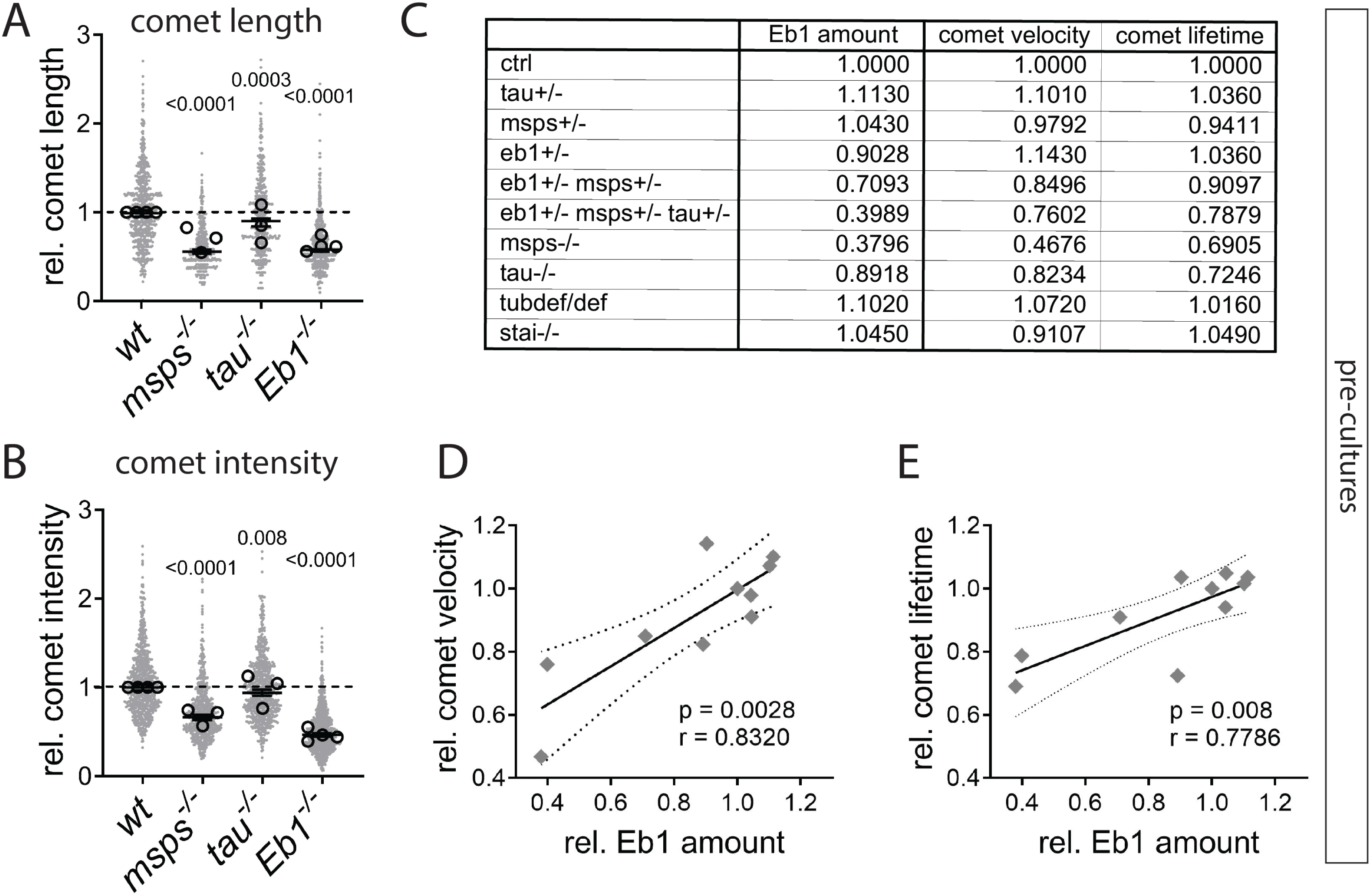
Correlation of different Eb1 comet properties. **A,B**) Eb1 amount at comets is calculated as the product of comet length (A) and the fluorescent mean intensity of Eb1 comets (B), which are both affected to similar degrees by homozygous condition of *msps^A^*, *tau^KO^* and *Eb1^04524^* in embryo-derived neurons cultured for 12hrs following 6 day pre-culture (6d pre); data were normalised to controls (dashed horizontal line) and are shown as scatter dot plot with median ± 95% confidence interval of at least three experimental repeats, large open circles in graphs indicate median/mean of independent biological repeats. P-values listed above each plot were obtained with Kruskall-Wallis ANOVA tests. **C**) The table lists data for Eb1 amounts (fixed neurons; compare Fig.1E-H, L) or for comet velocity/lifetime (live imaging; compare Fig.1M,N), all obtained from pre-cultured embryonic primary neurons carrying the same combinations of mutant alleles in homozygosis (indicated on the left; used alleles: *tub84B^def^*, *msps^A^*, *tau^KO/Df^*, *eb1^04524^*, *stai^KO^*). **D,E**) Plotting comet velocity or lifetime against Eb1 amounts from different genetic conditions shows good correlation (r and p-value determined via non-parametric Spearman correlation analysis). For raw data see Tab. T1-S2.

**Fig. 2-S1.**
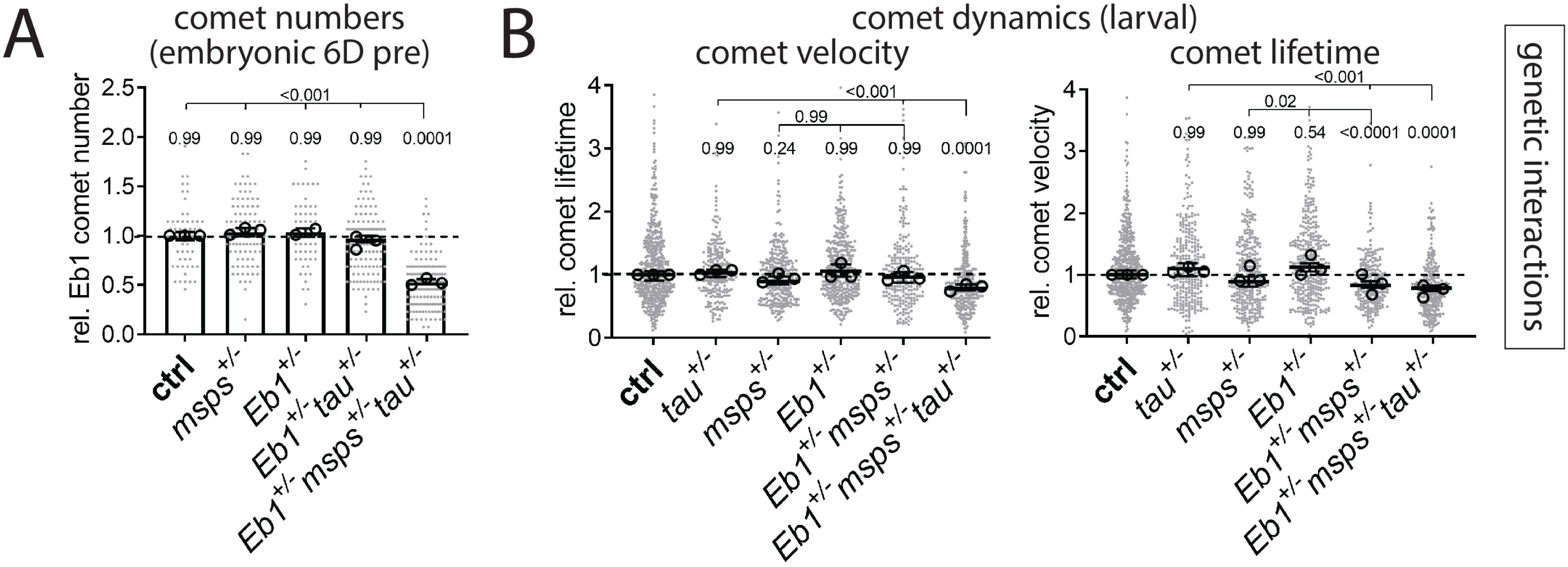
Comet numbers and dynamics in (trans-) heterozygous conditions **A**) Eb1 comet numbers in fixed primary neurons cultured for 12hrs following 5 day pre-culture. **B**) Comet velocity and lifetime obtained from live analyses of primary neurons cultured for 18hrs from CNSs of late larvae carrying single-, double- or triple-heterozygous conditions (complementing data in Fig. 2A). In all graphs, data were normalised to parallel controls (dashed horizontal lines) and are shown as scatter dot plots with median ± 95% confidence interval (B) or bar chart with mean ± SEM (A) of at least two experimental repeats; large open circles in graphs indicate median/mean of independent biological repeats. P-values obtained with Kruskal-Wallis ANOVA test are given above data points/bars; used alleles: *msps^A^*, *tau^KO^*, *Eb1^04524^*. For raw data see Tab. T2-S1.

**Fig. 2-S2.**
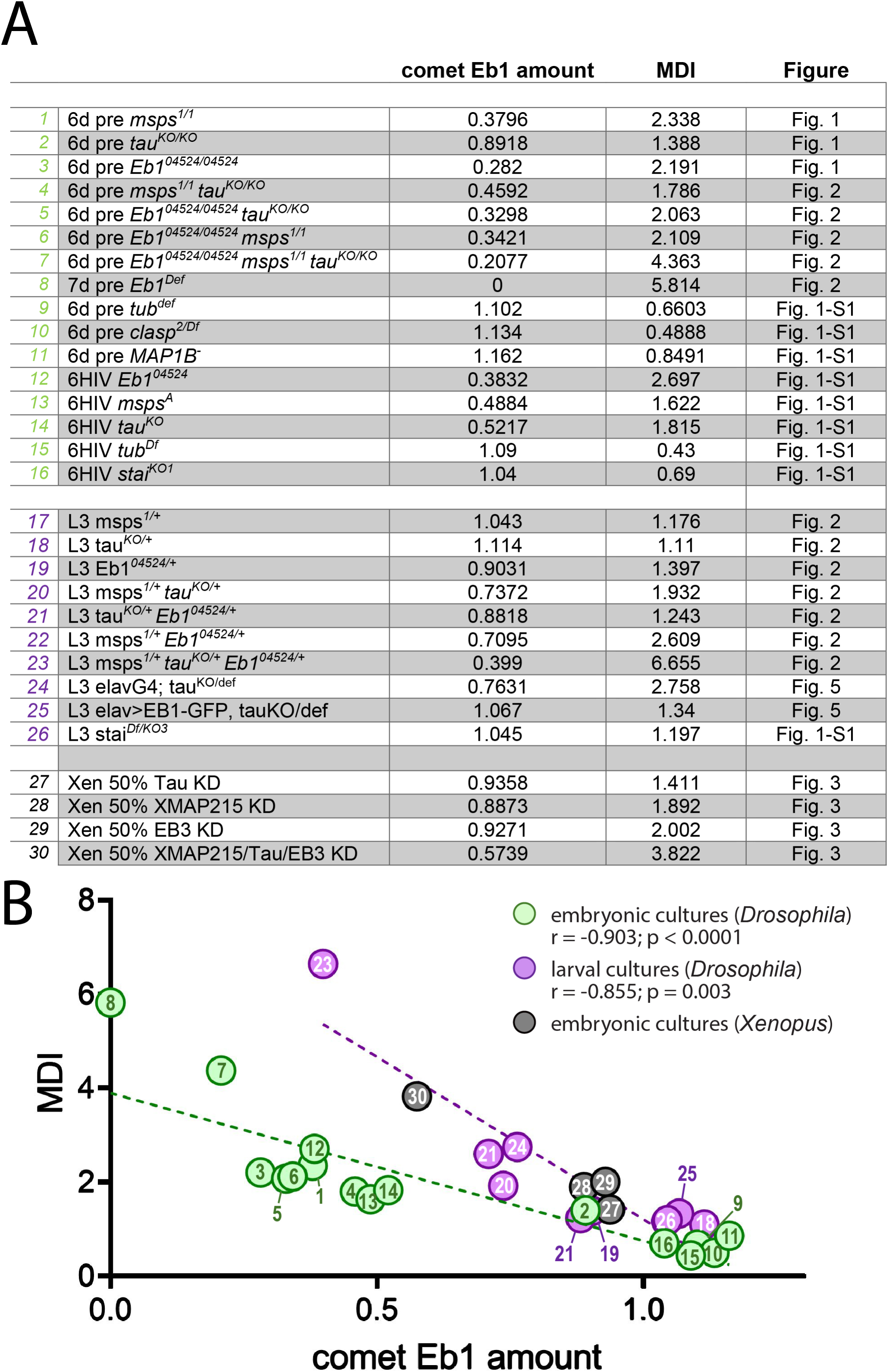
Increased Eb1 amounts correlate with decreased MT curling. The data and graph show further details behind the correlations displayed in Fig.2C. **A**) The table provides descriptions, data and references for the graph shown in B: coloured numbers (1^st^ column) correspond to numbers of data points in B; the different allelic combinations and culture conditions used (2^nd^ column) comprise embryonic neurons cultures for 6 hrs (6HIV), embryonic neurons cultured for 12 hrs following 6 day preculture (6d pre), neurons cultured from larval CNSs for 18hrs (L3); respective data for Eb1 amounts (3^rd^ column) and MT curling (4^th^ column) were obtained from different sets of experiments throughout this work (5^th^ column lists the figures from where these data originate). **B**) Correlation plot of the data shown in A, with numbers and colours of data points corresponding to the 1^st^ column; r and p-value determined via non-parametric Spearman correlation analysis. For raw data see Tab. T2-S2.

**Fig. 3-S1.**
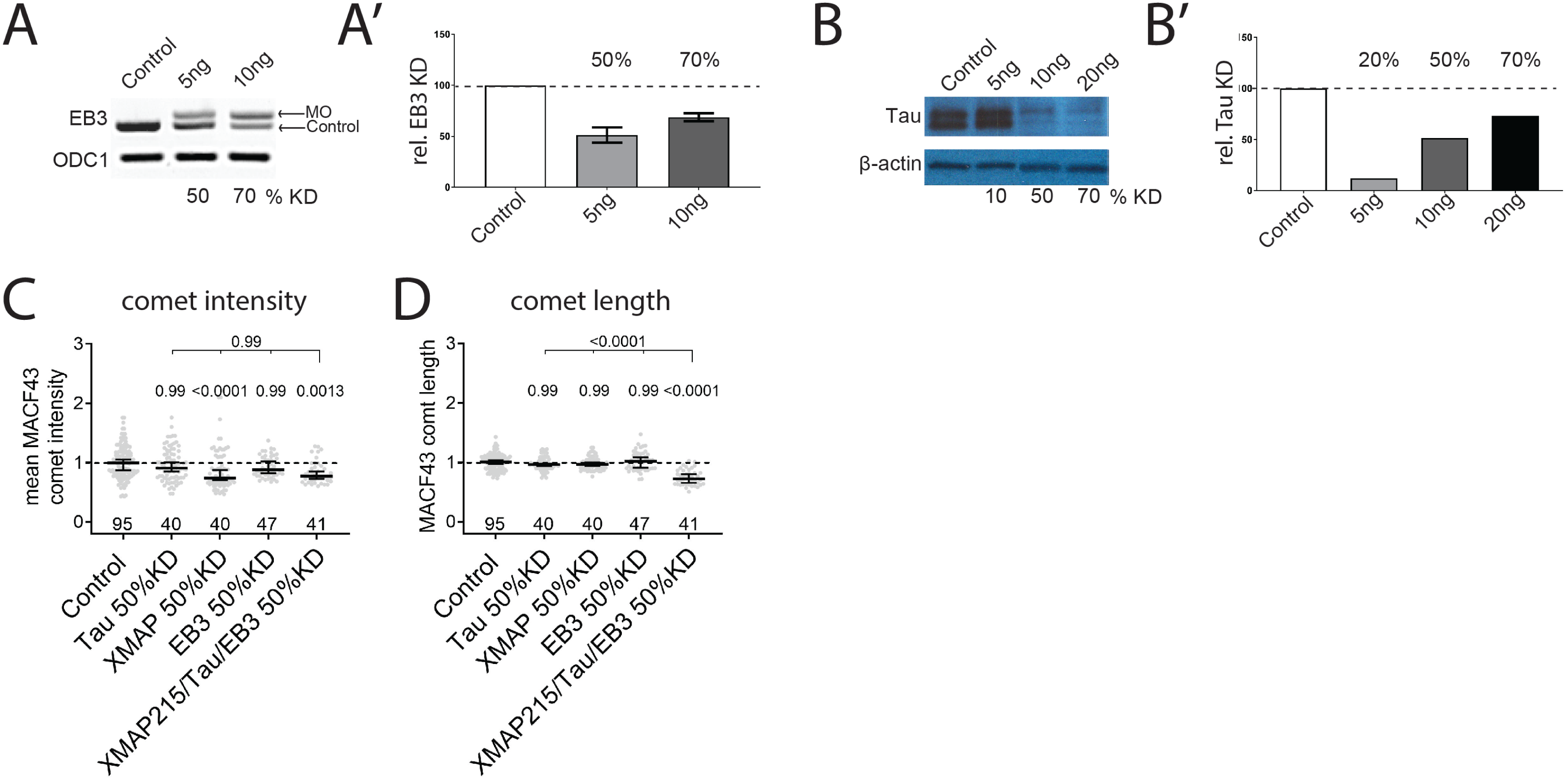
Support data for *Xenopus* experiments. **A-B’**) A RT-PCR DNA gel (A) and Western blot (B) and their quantifications (A’, B’) show the degrees of EB3/Tau knock-downs upon application of different morpholino concentrations (indicated on top in blots and at the bottom in graphs); ODC1 and ß-actin are used as loading controls; data are normalised to no-morpholino controls from two experimental repeats (dashed lines). 50% knock-down of XMAP215 was achieved by injecting 6 ng of the validated XMAP215 MO as described previously (Lowery *et al*., 2013; Slater *et al*., 2019). **C-D**) Different properties of MACF43::GFP comets (as indicated upon graphs) obtained from *Xenopus* primary neurons, either upon 50%; data were normalised to parallel controls (dashed horizontal lines) and are shown as median ± 95% confidence interval; merged sample numbers from at least two experimental repeats are shown at the bottom, P-values obtained with Kruskall-Wallis ANOVA and Dunn’s posthoc tests above data points. For raw data see Tab. T3-S1.

**Fig. 4-S1.**
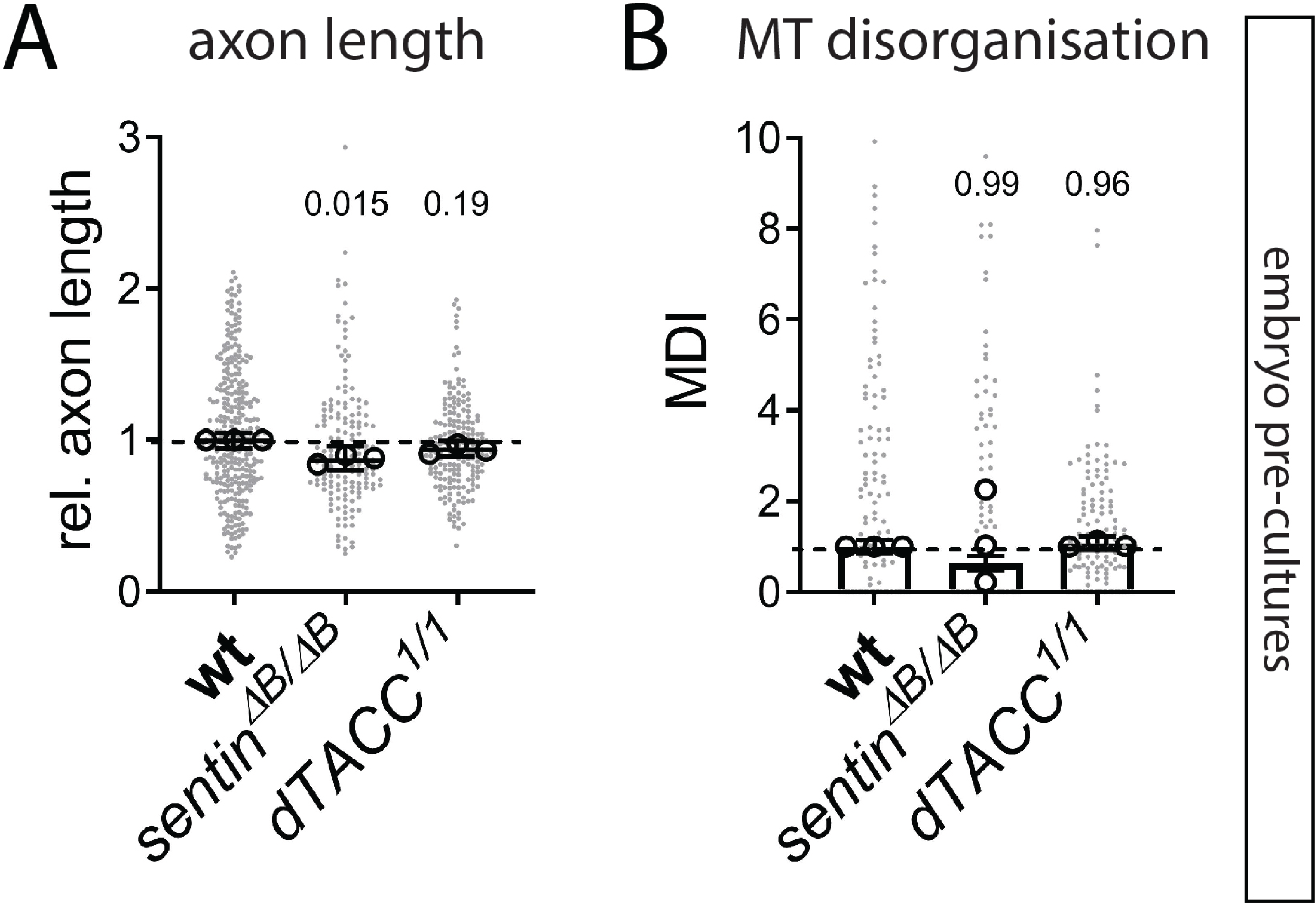
Loss of TACC or Sentin does not cause obvious axonal phenotypes. Axon length (**A**) and MT curling (**B**) embryonic 6d pre-cultured neurons which were either wild-type (wt) or homozygous mutant for *sentin* or *dTACC* (as indicated); data were normalised to parallel controls (dashed horizontal lines) and are shown as scatter dot plots with median ± 95% confidence interval (A) or mean ± SEM (B) from at least two experimental repeats; large open circles in graphs indicate median/mean of independent biological repeats. P-values obtained with Kruskal-Wallis ANOVA tests are shown above data points/bars. For raw data see Tab. T4-S1.

**Fig. 5-S1.**
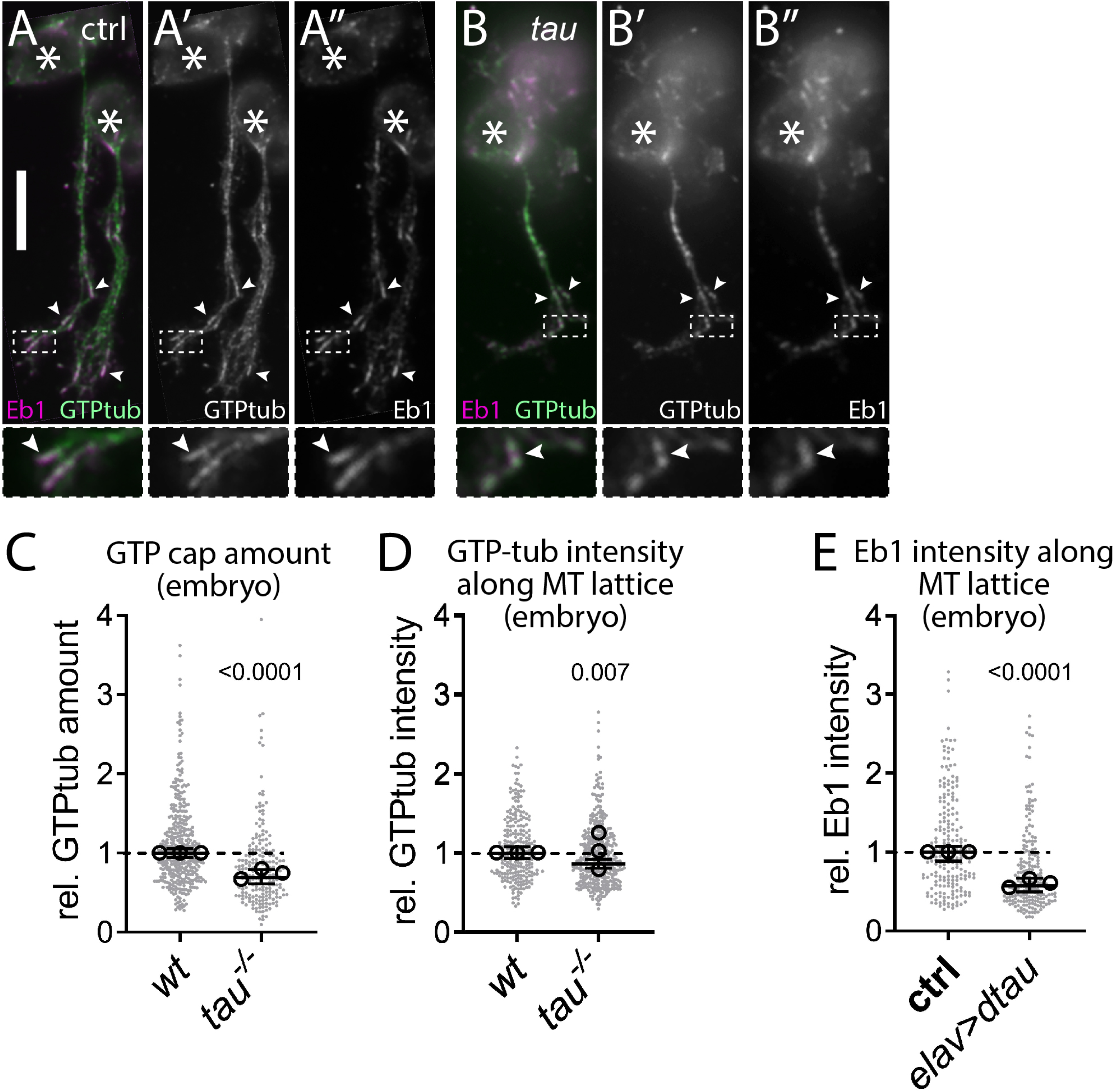
In Tau-deficient axons, GTP-tubulin is reduced at MT plus-ends but unchanged at lattices. **A-B’’**) Fixed primary neurons stained for Eb1 (magenta) and GTP-tubulin (green; the scale bar represents 10µm); asterisks indicate somata, dashed boxes the positions of the 4-fold magnified close-ups shown at the bottom, arrowheads point at Eb1::GFP comets and GTP caps. **C,D**) Graphs showing staining intensity of GTP-tubulin at MT plus-ends (C) and along the MT lattice (D) of embryonic neuron at 6 HIV. **E)** Graphs showing staining intensity of Eb1 along the MT lattice of neurons without/with *elav-Gal4*-driven expression of *Drosophila* Tau (*dtau*). Overexpression of *dtau* leads to a reduction of Eb1 at the MT shaft; data were normalised to parallel controls (dashed horizontal lines) and are shown as scatter dot plots with median ± 95% confidence interval from at least two independent repeats with 3 experimental replicates; large open circles in graphs indicate median/mean of independent biological repeats. P-values obtained with Kruskall-Wallis ANOVA test above data points/bars. For raw data see Tab. T5-S1.

**Fig. 5-S2.**
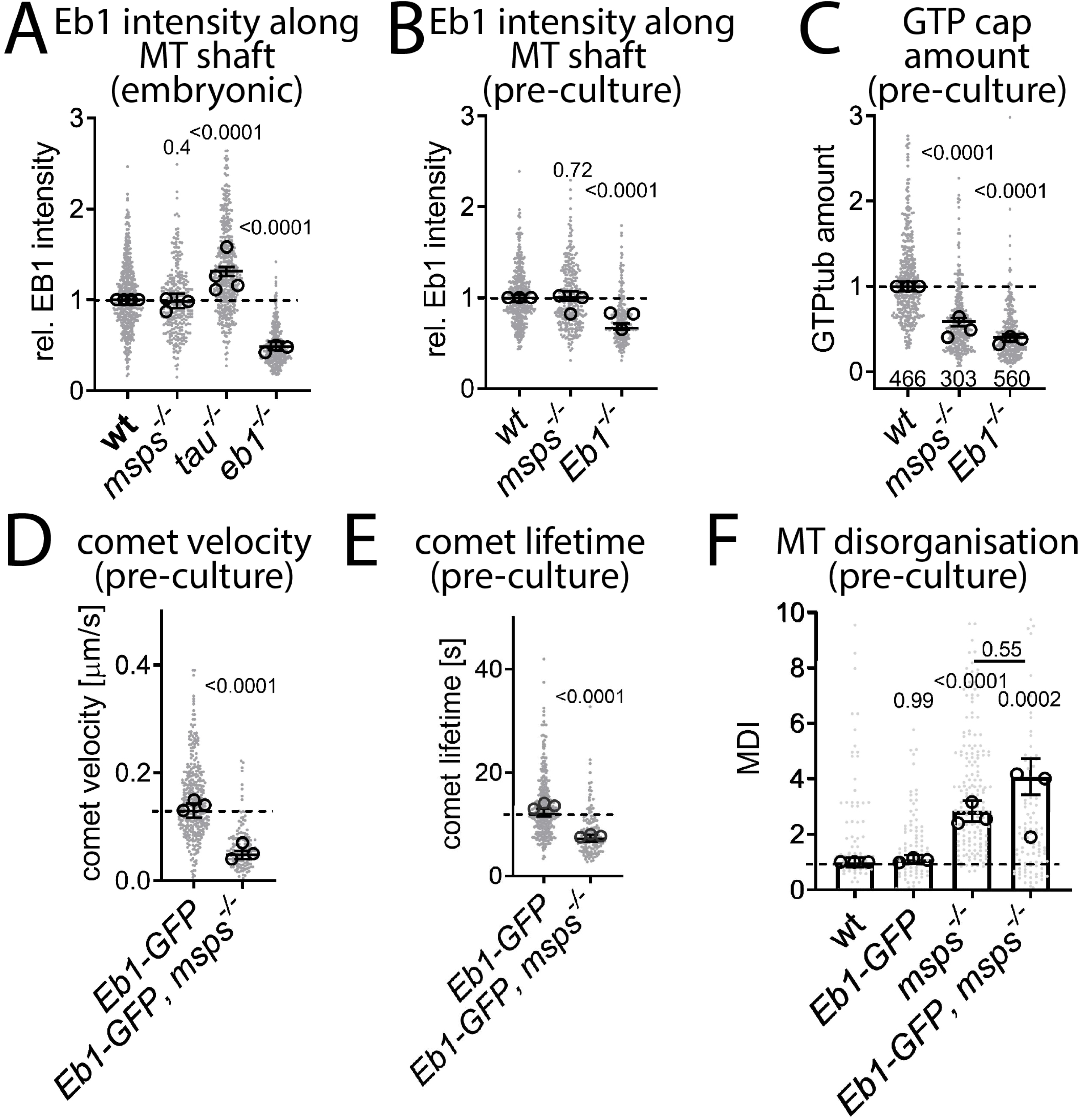
Details of Eb1-Msps cross-regulation. **A,B**) Eb1 lattice localisation is unaffected by loss of Msps in primary neurons at 6HIV (A) or at ∼12HIV following 6 day pre-culture (B). **C**) Staining intensity of GTP-tubulin at MT plus-ends is reduced in *msps^A/A^* and *Eb1^04524/04524^* mutant embryonic neurons at ∼12 HIV following 6 day pre-culture. **D-F**) Expressing Eb1::GFP via *elav-Gal4* does not rescue comet velocities (D), comet lifetime (E) and MT curling (F) in primary neurons of *msps^A/146^* mutants (cultured12 HIV following 6 day pre-culture); data were normalised to parallel controls (dashed horizontal lines) and are shown as scatter dot plots with mean ± SEM (F) or median ± 95% confidence interval (A-E) from at least two experimental repeats; large open circles in graphs indicate median/mean of independent biological repeats. P-values obtained with Kruskal-Wallis ANOVA tests and Dunn’s posthoc analyses are shown above data points/bars. For raw data see Tab. T5-S2

**Movie M1**. Msps::GFP and Eb1::RFP jointly track MT plus-ends. Live movie of a wild-type neuron co-expressing Msps::GFP and Eb1::RFP; for stills see Fig.4A-A’’. As indicated, single channels are shown on the left and middle, and the combined movie on the right. The movie was acquired at 0.5 frames per second and plays at 0.5 s per frame. The scale bar indicates 10 μm.

**Movie M2**. Msps plus-end localisation in wild-type neurons. Live movie of a wild-type neuron expressing Msps::GFP; for a still see Fig.4C. The movie was acquired at 1 frame per second and plays at 0.2 s per frame. The scale bar indicates 10 μm.

**Movie M3**. Msps plus-end localisation is impaired in the absence of Eb1. Live movie of an *Eb1^04524/04524^* mutant neuron expressing Msps::GFP; for a still see Fig.4C’. The movie was acquired at 1 frames per second and plays at 0.2 s per frame. The scale bar indicates 10 μm.

